# A *C. elegans* Zona Pellucida domain protein functions via its ZPc domain

**DOI:** 10.1101/2020.05.25.115105

**Authors:** Jennifer D. Cohen, Jessica G. Bermudez, Matthew C. Good, Meera V. Sundaram

## Abstract

Zona Pellucida domain (ZP) proteins are critical components of the body’s external-most protective layers, apical extracellular matrices (aECMs). Although their loss or dysfunction is associated with many diseases, it remains unclear how ZP proteins assemble in aECMs. Current models suggest that ZP proteins polymerize via their ZPn subdomains, while ZPc subdomains modulate ZPn behavior. Using the model organism *C. elegans*, we investigated the aECM assembly of one ZP protein, LET-653, which shapes several tubes. Contrary to prevailing models, we find that LET-653 localizes and functions via its ZPc domain. Furthermore, the ZPc domain is inhibited by the ZPn domain and cleavage of the LET-653 C-terminus relieves this inhibition. *In vitro*, the ZPc, but not ZPn, domain formed crystalline aggregates. These data offer a new model for ZP function whereby the ZPc domain is primarily responsible for matrix incorporation and tissue shaping.

## Introduction

The apical extracellular matrices (aECMs) that line the insides, or lumens, of biological tubes are critical for shaping and maintaining tube structure (Luschnig & Uv, 2014). For example, the aECM of the blood vasculature, the glycocalyx, prevents capillary loss (Gaudette, Hughes, & Boller, 2020). The surfactant that lines the lung’s narrow tubes decreases surface tension to prevent alveoli collapse (Halliday, 2017). Loss or dysfunction of aECMs are associated with a number of human diseases (Gaudette et al., 2020; Gordts & Esko, 2018; Halliday, 2017; Hansson, 2019).

Despite aECMs’ importance in human health, we know little about aECM assembly. aECMs are complex meshworks of glycoproteins, glycosaminoglycans, proteoglycans, lipids, and lipoproteins (Gaudette et al., 2020; Gordts & Esko, 2018; Halliday, 2017; Hansson, 2019; Luschnig & Uv, 2014). Individual aECM components can sometimes form insoluble fibrils or water-trapping gels *in vitro* (Husain et al., 2006; Hwang, Olson, Esko, & Horvitz, 2003; Jovine, Qi, Williams, Litscher, & Wassarman, 2002; Lane, Koehl, Wilt, & Keller, 1993; Porter & Tamm, 1955; Syed et al., 2012), but how different components come together *in vivo* is more challenging to decipher. Luminal aECMs are usually difficult to visualize *in vivo* because they are translucent and their structures can be destroyed under desiccating fixation conditions (Chappell et al., 2009). Where aECMs have been observed, they often form multi-layered structures (Chappell et al., 2009; Johansson, Sjovall, & Hansson, 2013). For example, the vascular aECM may be up to several microns thick and contains at least three layers with distinct electron density (Chappell et al., 2009; Gaudette et al., 2020). Generating an aECM requires inhibiting assembly of its various components until after secretion, and then encouraging assembly with appropriate partners at the right time and place. Defects in any of these regulatory steps could disrupt aECM organization and contribute to disease.

Zona Pellucida domain, or ZP, proteins are a widespread and abundant aECM protein family (Plaza, Chanut-Delalande, Fernandes, Wassarman, & Payre, 2010). ZP proteins were originally identified as the major components of the mammalian oocyte’s extracellular coat, also called the Zona Pellucida (Gupta, 2018). ZP proteins were later discovered within many somatic aECMs, including those of the kidney tubules (Umod (Porter & Tamm, 1955), Oit3 (Yan, Zhang, Huang, Shen, & Han, 2012)), inner ear (Tecta, Tectb (Legan, Rau, Keen, & Richardson, 1997)), gut (GP2 (Op De Beeck, Vermeire, Rutgeerts, & Bossuyt, 2012), DMBT3 (Renner et al., 2007)) and vasculature (Endoglin (McAllister et al., 1994), Betaglycan (Wong, Hamel, Chevalier, & Philip, 2000)). Loss or dysfunction of these ZP proteins is associated with many human diseases, including chronic kidney disease, deafness, and hereditary hemorrhagic telangiectasia (HHT) (Cummings et al., 2018; Mapes et al., 2017; McAllister et al., 1994; Mustapha et al., 1999; Renner et al., 2007; Roggenbuck et al., 2009; Verhoeven et al., 1998; Wheeler & Thomas, 2019; Wong et al., 2000). Several ZP proteins are thought to act primarily by trapping ligands and affecting signaling processes (Lin, Hu, Zhu, Woodruff, & Jardetzky, 2011; Saito et al., 2017), but many form fibrils or aggregates that may play structural roles within the aECM (Bokhove & Jovine, 2018; Jovine et al., 2002; Porter & Tamm, 1955; Shen et al., 2009). How ZP proteins assemble into complex extracellular matrix structures *in vivo* is poorly understood.

Although originally described as a single domain, ZP domains are now recognized to contain two separate domains, ZPn and ZPc, that are usually, but not always, found in tandem (Wilburn & Swanson, 2017). Crystal structures reveal that ZPn and ZPc domains independently take on Immunoglobulin (Ig)-like folds held in place by disulfide bonds between paired cysteines (Han et al., 2010; Lin et al., 2011; Saito et al., 2017). ZPn domains typically have four evenly spaced cysteines, while ZPc domains can have four, six, or eight cysteines with some variability in spacing (Bokhove & Jovine, 2018; Cohen, Flatt, Schroeder, & Sundaram, 2019). ZPc domains are usually followed by a furin-type consensus cleavage site (CCS), then a hydrophobic C-terminus, and sometimes a transmembrane domain or GPI anchor.

Several ZPn domains can polymerize spontaneously to form fibrils *in vitro*, but functions for ZPc domains are less well defined (Bokhove et al., 2016; Darie, Janssen, Litscher, & Wassarman, 2008; Jovine, Janssen, Litscher, & Wassarman, 2006; Litscher, Janssen, Darie, & Wassarman, 2008; Louros et al., 2016; Louros, Iconomidou, Giannelou, & Hamodrakas, 2013). Studies of two non-polymerizing ZP proteins, Endoglin and Betaglycan, suggest that ZPc domains can dimerize and may mediate other protein-protein interactions (Lin et al., 2011; Saito et al., 2017). However, due to challenges in expressing ZPc domains in cell culture (Bokhove et al., 2016; Monne, Han, Schwend, Burendahl, & Jovine, 2008), the roles or polymerization capabilities of ZPc domains independent of ZPn domain have not been assessed.

A prevailing model has been that ZPc domains serve as negative regulators of ZPn polymerization. *In vivo*, an external hydrophobic patch (EHP) C-terminal to the CCS is generally required for ZP protein secretion (Jimenez-Movilla & Dean, 2011; Jovine, Qi, Williams, Litscher, & Wassarman, 2004; Williams & Wassarman, 2001), and cleavage at the CCS occurs prior to matrix incorporation (Boja, Hoodbhoy, Fales, & Dean, 2003; Litscher, Qi, & Wassarman, 1999). Structural data showed that the EHP is part of the ZPc fold (Han et al., 2010), and suggested a model in which ZPn polymerization is inhibited prior to secretion via an intramolecular interaction with the ZPc domain and the C-terminus (Bokhove et al., 2016; Jovine et al., 2004). This interaction may be stabilized by binding between the EHP and an internal hydrophobic patch (IHP) in the linker between the ZPn and ZPc domains (Jovine et al., 2004; Zhao et al., 2003). Under this model, cleavage at the CCS evicts the EHP and C-terminus, releases ZPc-mediated inhibition, and enables ZPn polymerization (Brunati et al., 2015; Jovine et al., 2004). However, this model is based on *in vitro* studies of only a few proteins, most of which were modified to promote efficient secretion (Bokhove & Jovine, 2018; Fahrenkamp, Algarra, & Jovine, 2020). It remains unclear whether the ZPc and C-terminus have roles other than facilitating ZPn secretion and assembly, and if and how the ZPn and ZPc domains impact one another’s function *in vivo*.

ZP proteins are important structural components of cuticle and other aECMs in invertebrates, including *Caenorhabditis elegans* and *Drosophila* (Fernandes et al., 2010; Plaza et al., 2010). *C. elegans* has an unusually large set of ZP proteins (Cohen et al., 2019), many of which affect shaping of specific epithelia (Forman-Rubinsky, Cohen, & Sundaram, 2017; Gill et al., 2016; Kelley et al., 2015; Low et al., 2019; Muriel et al., 2003; Sapio, Hilliard, Cermola, Favre, & Bazzicalupo, 2005; Sebastiano, Lassandro, & Bazzicalupo, 1991; Vuong-Brender, Suman, & Labouesse, 2017; Yu, Nguyen, Hall, & Chow, 2000). Furthermore, *C. elegans* is genetically tractable and is transparent, allowing live imaging of the aECM (Corsi, Wightman, & Chalfie, 2015). These features make *C. elegans* a particularly attractive model for studying ZP structure/function relationships and matrix assembly.

Here, we investigate the matrix assembly of the *C. elegans* ZP protein LET-653, which shapes multiple tube types. We identify the ZPc domain as the critical domain for LET-653 function and matrix localization and discover that cleavage releases inhibition of the ZPc domain by the ZPn domain. We also show for the first time that an isolated ZPc domain promotes formation of *in vitro* aggregates. Together, these data offer a novel model for ZP protein function and matrix assembly.

## Results

### Background: LET-653 is a luminal ZP protein that shapes the duct and vulva tubes

LET-653 (Figure 1A) is a component of a transient pre-cuticular luminal matrix that shapes multiple developing tubes (Cohen et al., 2020; Gill et al., 2016). LET-653 is essential to shape the tiny excretory duct tube, a unicellular tube that is less than a micron in diameter (Gill et al., 2016). All *let-653* mutants die as young larvae due to discontinuities in this tube lumen that prevent fluid excretion (Figure 1B) (Buechner, Hall, Bhatt, & Hedgecock, 1999; Gill et al., 2016). *let-653* mutants rescued in the duct cell survive into adulthood, but show more subtle defects in the shaping of other epithelial tissues, including the vulva, which is a large multicellular tube that, in adulthood, serves as the passageway through which embryos are laid (Forman-Rubinsky et al., 2017; Gill et al., 2016) (Cohen et al., 2020; Gill et al., 2016) (Figure 1E). The excretory duct tube provides the most sensitive assay for LET-653 function, while both the duct (Figure 1C-D) and the vulva (Figure 1E-G) allow rapid visual assessment of matrix incorporation via imaging of fluorescently-tagged LET-653 proteins in live animals.

**Figure 1.**
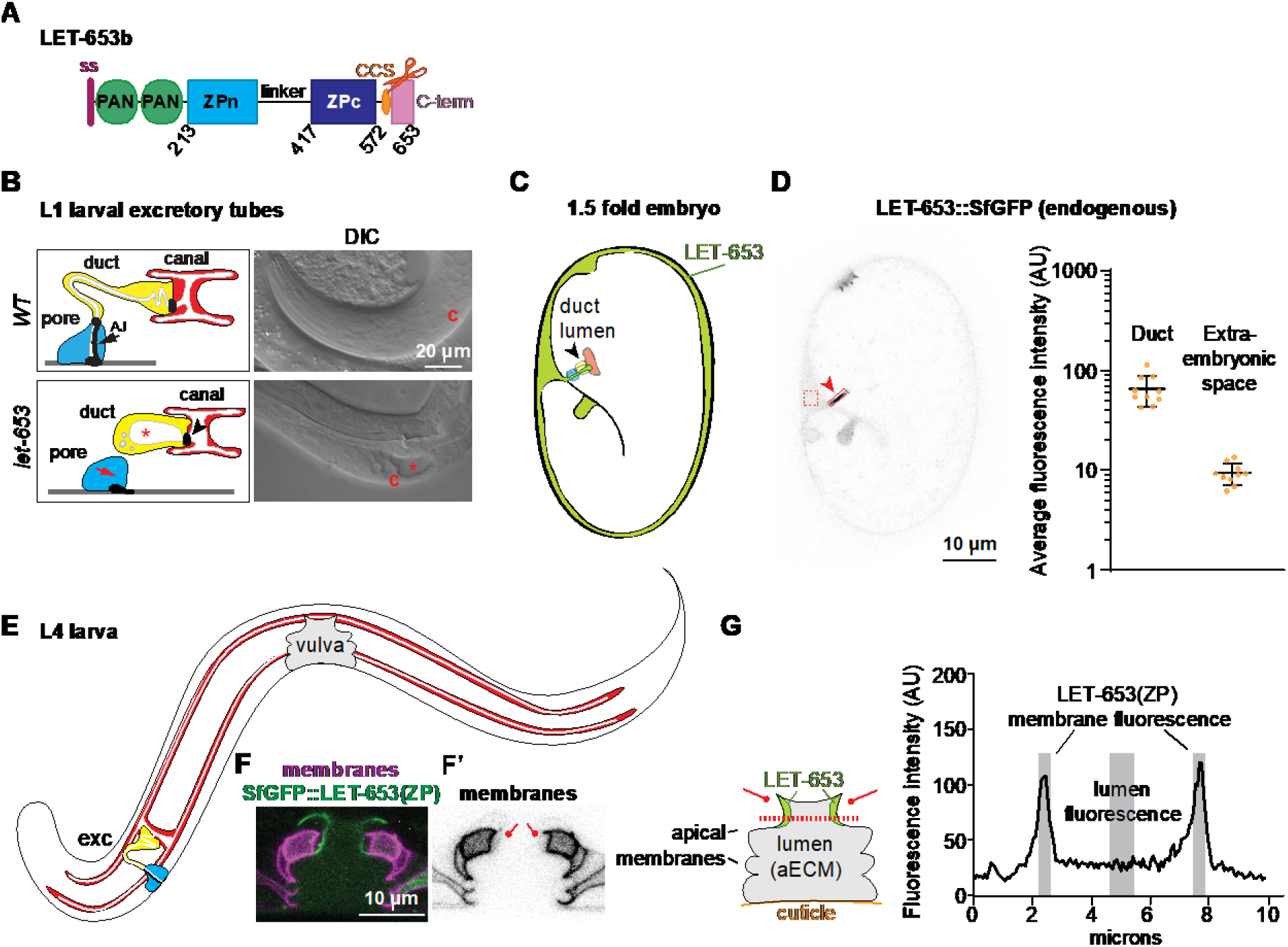
LET-653 is a luminal ZP protein that shapes *C. elegans* tubes. A) Diagram of LET-653 isoform b, on which all structure/function constructs were based. LET-653 has an N-terminal signal peptide (ss, pink) but lacks any other membrane-association domain. PAN; PAN/Apple domains, which are also found in the blood clotting factor plasminogen (Tordai, Banyai, & Patthy, 1999). CCS (orange oval and scissors); consensus furin-type cleavage site (RRYR) located after the ZP domain and before the C-terminus. The linker is predicted to be N-glycosylated and heavily O-glycosylated. Pink box represents C-terminus. The LET-653 ZP domain aligns well with other ZP proteins, but has several unusual features. The ZPn domain lacks a key tyrosine residue thought to be important for ZPn polymerization (Gill et al., 2016; Monne et al., 2008), and the linker region between the ZP subdomains is unusually long, a feature whose impact is unclear (Cohen et al., 2019). B) Cartoons of the duct and pore cells of the excretory system and DIC images of *let-653* mutants and *WT C. elegans* at L1 stage. Yellow; duct cell. Blue; pore cell. White; lumen. Black; cell junctions. Asterisk; luminal dilation. c; canal cell nucleus. B) Diagram of *C. elegans* 1.5 fold embryo. White; embryo. Gray; extraembryonic space. Arrowhead indicates duct/pore lumen. Red; canal cell. Yellow; duct cell. Blue; pore cell. D) Confocal image of a 1.5 fold embryo with CRISPR-tagged full-length LET-653::SfGFP (Cohen et al., 2020) with quantification of duct and extraembryonic fluorescence intensity. LET-653 is enriched in the duct lumen (arrowhead), buccal cavity and rectum. Fluorescence intensity was calculated within the indicated boxes to estimate matrix incorporation and overall expression levels, respectively. SfGFP alone shows minimal duct lumen fluorescence but strong extraembryonic fluorescence (Gill et al., 2016), while endogenously tagged full-length LET-653 shows strong duct lumen fluorescence but minimal extraembryonic fluorescence (Cohen et al., 2020). Quantification of average endogenous LET-653 fluorescence intensity per pixel in boxes drawn within the duct lumen and extraembryonic space. n = 10. AU; Arbitrary units. Error bars represent standard error. E) Cartoon of L4 larva. F-F’) Confocal images of the vulva apical membranes, marked by PH::mCherry, and LET-653(ZP), which lines the dorsal vulE and vulF membranes. F’) vulE and vulF membranes are faintly labelled by PH::mCherry (lines). G) Cartoon of vulva lumen and quantification of LET-653 fluorescence. The lumen is lined by 22 cells whose apical membranes define lumen shape. Cuticle lines the base of the vulva and a membranous hymen covers the top, generating a closed tube. Drawing a line across the dorsal region of the vulva allows quantification of LET-653 membrane vs. luminal localization. LET-653 fluorescence across the dashed line shows two peaks near the apical membranes and decreased fluorescence in the center of the lumen. Black; LET-653 fluorescence. Gray; membrane-associated and luinal regions whose fluorescence intensities were averaged. A line of 10 pixels thick (0.63 microns) was drawn across the vulva in the position indicated in panel H. Fluorescence intensity from 5 points proximal to each apical membrane (10 values total) were averaged and divided by the average fluorescence intensity across 10 points from the center of the vulva lumen. AU; arbitrary units.

Previously, we demonstrated that the LET-653 ZP domain is responsible for both tube shaping and incorporation into a membrane anchored aECM (Cohen et al., 2020; Gill et al., 2016). LET-653 is a secreted protein with two N-terminal PAN/Apple (plasminogen) domains and a C-terminal ZP domain (Jones & Baillie, 1995) (Figure 1A). The PAN and ZP domains each confer different patterns of matrix localization, but only the ZP domain is capable of rescuing *let-653* mutant lethality and both duct and vulva tube shape defects (Cohen et al., 2020; Gill et al., 2016). As observed for most other ZP proteins, the LET-653 C-terminus is cleaved, though LET-653 has no transmembrane or membrane association domain (Gill et al., 2016). The cleaved C-terminus stays associated with the rest of the ZP domain *in vivo*, such that tagging LET-653(ZP) at either the N- or C-terminus yields the same localization pattern (Gill et al., 2016). Fluorescence recovery after photobleaching (FRAP) analyses indicated that LET-653(ZP) has low mobility, consistent with matrix incorporation (Gill et al., 2016).

### LET-653 functions via the ZPc domain

Transgenic structure/function experiments were performed to dissect ZP domain aECM assembly and tube shaping function. ZP domain fragments were expressed under a 2.2 kb *let-653* promoter fragment and tagged with Superfolder GFP (SfGFP) (Gill et al., 2016; Pedelacq, Cabantous, Tran, Terwilliger, & Waldo, 2006). These transgenes were expressed at levels comparable to endogenous, SfGFP-tagged LET-653 as assessed by fluorescence levels (Figure 1D, S1). For each transgene, multiple independent lines were tested in three assays. To determine function, we tested the ability of transgenes to restore viability to *let-653(cs178)* null mutants (Figure 1B). To assay localization, we imaged the developing duct tube in 1.5 fold embryos (Figure 1C-D), and the developing vulva tube in L4 larvae (Figure 1F-G). In such assays, the positive control LET-653(ZP) localized transiently to the excretory duct lumen (Figures 2A, S2) and along specific apical membranes within the vulva tube (Figure 1F-G, 2A).

**Figure 2.**
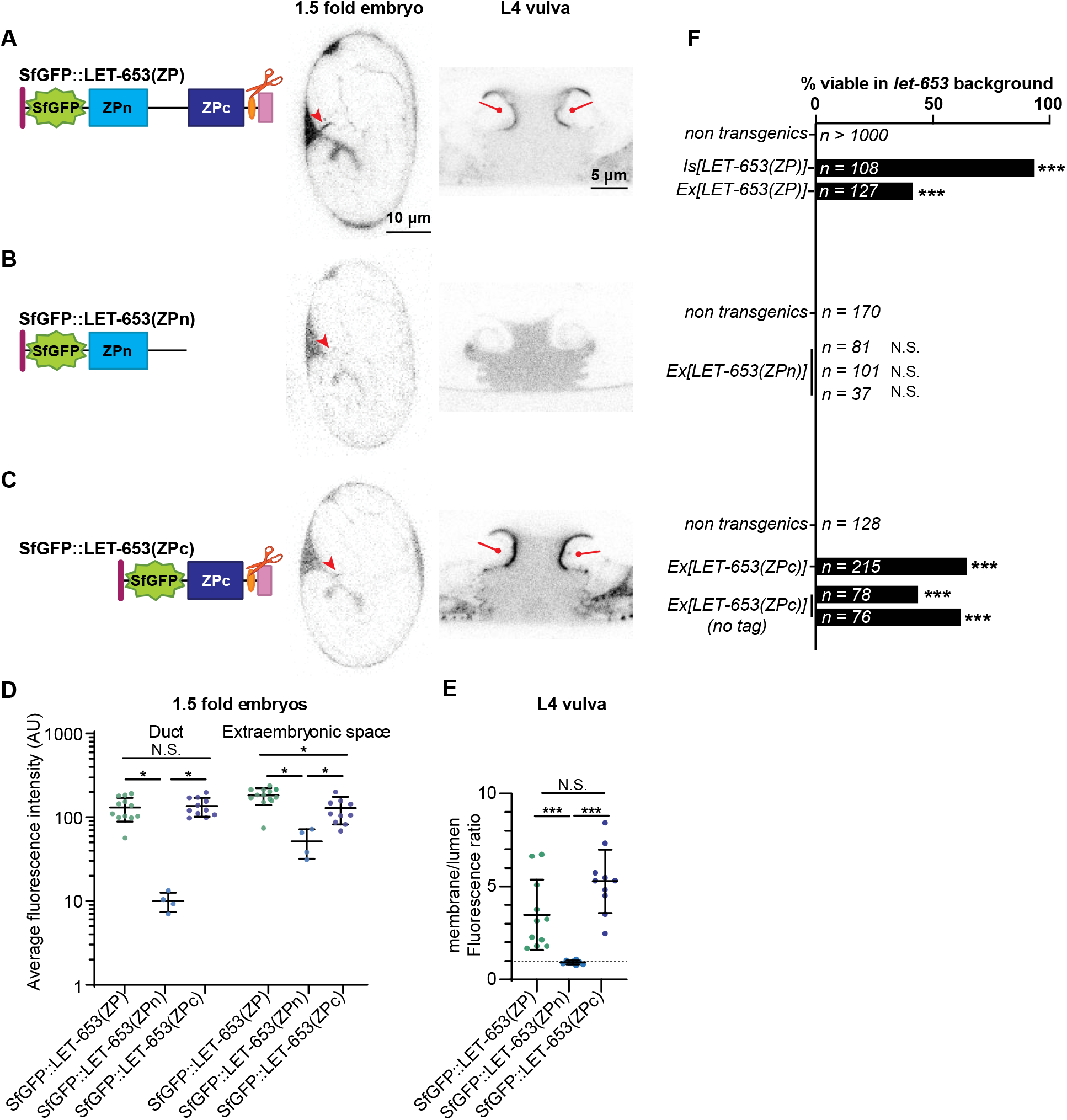
The ZPc domain is necessary and sufficient for LET-653 function. Here and in all subsequent figures, cartoons depict the LET-653 fragments tested and the location of the SfGFP tag (green). A) Full ZP domain. SfGFP::LET-653(ZP) is consistently visible in the duct lumen of 1.5-fold embryos (arrowhead) and along the dorsal apical membranes of the L4 vulva (lines). B) ZPn domain, including the linker. SfGFP::LET-653(ZPn) was occasionally observed in the 1.5 fold duct lumen, but never lined the vulva apical membranes. C) ZPc domain, including the C-terminus (Cterm). SfGFP::LET-653(ZPc) consistently lined the duct lumen and the vulva dorsal apical membranes (lines). D) Duct and extraembryonic fluorescence in 1.5 fold embryos were quantified as shown in Figure 1. Duct fluorescence levels were generally similar to levels of endogenous, full-length LET-653 (Figure 1D), although there were far greater levels of extraembryonic fluorescence, indicating overexpression and/or ectopic localization. p values generated using a two tailed Mann Whitney U test. Duct fluorescence: SfGFP::LET-653(ZP) vs SfGFP::LET-653(ZPc), p=0.7399. SfGFP::LET-653(ZP) vs SfGFP::LET-653(ZPn), *p=0.0011. SfGFP::LET-653(ZPn) vs SfGFP::LET-653(ZPc), *p=0.0015. Extraembryonic fluorescence: SfGFP::LET-653(ZP) vs SfGFP::LET-653(ZPc), *p=0.0086. SfGFP::LET-653(ZP) vs SfGFP::LET-653(ZPn), *p=0.0011. SfGFP::LET-653(ZPn) vs SfGFP::LET-653(ZPc), *p=0.0029. Error bars; standard error. E) Ratio of apical membrane-associated to lumen fluorescence in the L4 vulva (see Figure 1). ***p<0.0001, Mann Whitney two-tailed U test. Dashed line indicates a ratio of 1, where membrane fluorescence is equal to luminal fluorescence. Error bars; standard error. F) Both integrated (Is) and extrachromosomal (Ex) transgenes containing LET-653(ZP) rescued *let-653* mutant lethality. Non-transgenic animals are pooled siblings from experiments in each group; non-transgenic animals never survive. 0/3 independent transgenic lines containing LET-653(ZPn) were functional, but 3/3 lines containing LET-653(ZPc) (either with or without an SfGFP tag) were functional. ***p<0.0001, two-tailed Fisher’s Exact test.

To determine which part of the LET-653 ZP domain is functionally important, we first expressed constructs containing either the ZPn domain and linker or the ZPc domain and C-terminal tail (referred to henceforth as simply ZPn or ZPc). Both proteins were efficiently secreted, but LET-653(ZPn) was expressed at slightly lower levels than LET-653(ZPc) in embryos (Figure 2B-E). The ZPn domain transgene did not rescue *let-653* mutant lethality (Figure 2F). This protein localized weakly to the duct lumen and had no specific localization in the vulva (Figure 2B, D-E). In contrast, LET-653(ZPc) rescued *let-653* mutants (Figure 2F). This protein robustly lined the duct and vulva lumens (Figure 2C-E). These data indicate that LET-653(ZPc) is sufficient for both function and proper localization, and suggest that this domain binds to a partner that helps recruit it to the membrane-anchored matrix.

To test whether the observed function and localization were due to activity of the main part of the ZPc domain or to the C-terminal tail that follows the CCS, we attempted to express each segment independently. The LET-653 C-terminus contains a hydrophobic region that may correspond to an EHP (Figure 3A). To generate a truncated ZPc construct, we removed most (42/78 amino acids) of the C-terminus, leaving only this short, hydrophobic region (Figure 3A, B). LET-653(ZPc-½Cterm) was not secreted and did not rescue *let-653* mutant duct defects (Figure 3B, D). This is consistent with prior results showing that the C-terminus is required for secretion of LET-653(full ZP)-containing proteins, though not for secretion of LET-653(ZPn) ((Gill et al., 2016); Figure 2B). We conclude that the full C-terminus is required for proper trafficking and secretion of the ZPc domain.

**Figure 3.**
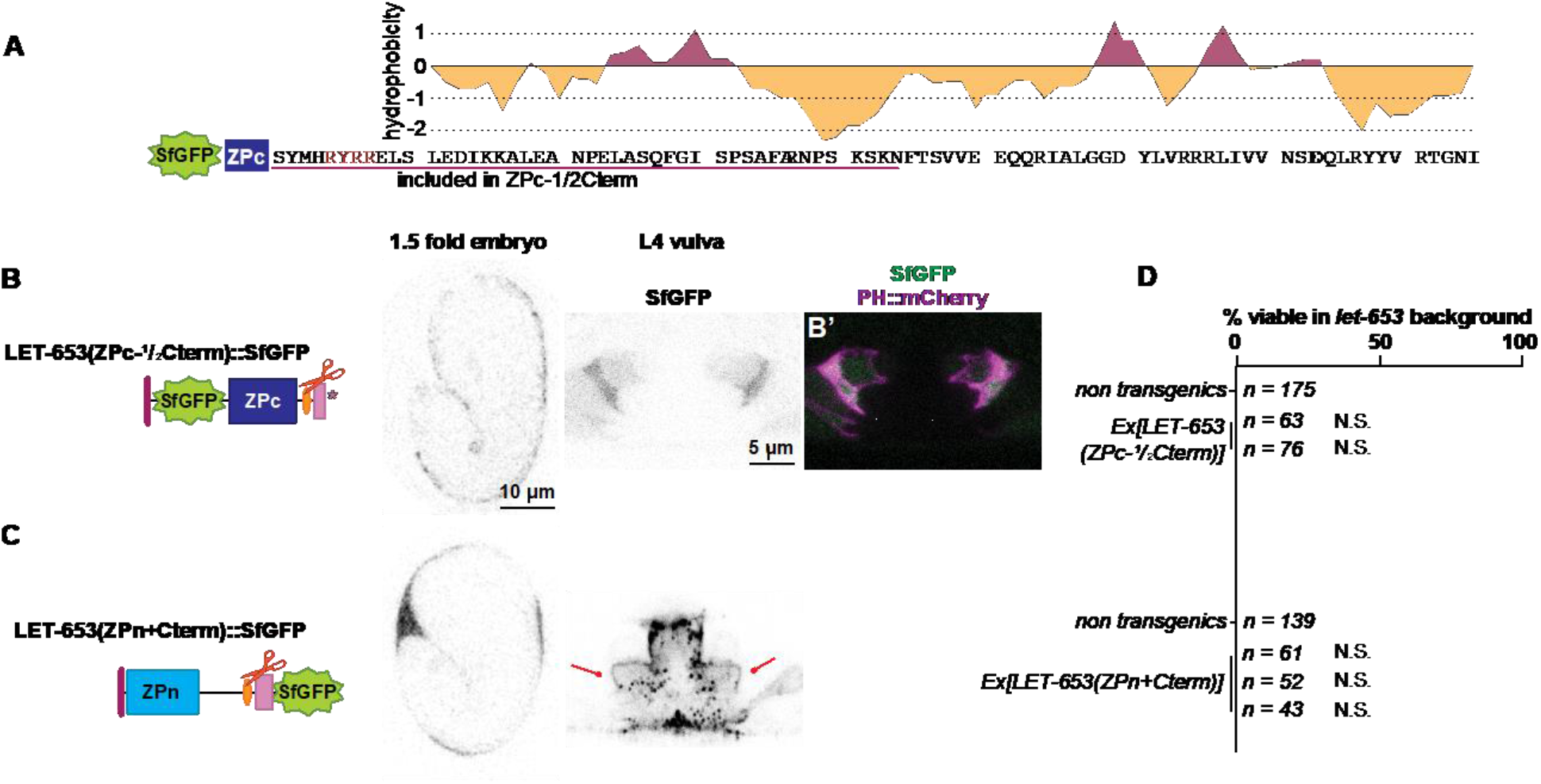
The LET-653 C-terminus confers aggregation activity but is not sufficient for function. A) Diagram of the LET-653 C-terminus. Cleavage site is shown in red. Potential external hydrophobic region (EHP); pink underline. Hydrophobicity plot generated using Kyte and Doolittle algorithm (Kyte & Doolittle, 1982). B) SfGFP::LET-653(ZPc-½Cterm) is truncated after the region underlined in pink in A. LET-653(ZPc-½Cterm) was not secreted. Absence of LET-653 from the duct, extraembryonic space, and vulva lumen was assessed visually via epifluorescence microscopy (n=20 each). B’) Overlay with SfGFP::LET-653(ZPc-½Cterm) and the membrane marker PH::mCherry, indicating all SfGFP::LET-653(ZPc-½Cterm) is enclosed in membrane. C) LET-653(ZPn+Cterm)::SfGFP. The entire C-terminus, starting at the end of the ZPc subdomain, was added to the end of the ZPn domain. Although efficiently secreted, LET-653(ZPn+Cterm) was rarely observed in the embryonic duct lumen (3/20 embryos) and had a punctate appearance in the vulva lumen (n=10). Some LET-653(ZPn+Cterm) lined vulva apical membranes (lines). As LET-653(ZPn+Cterm) had punctate localization, its enrichment at the vulva membrane vs. the center of the vulva lumen was not quantified. D) Two independent transgenes containing LET-653(ZPc-½Cterm), and three transgenes containing LET-653(ZPn+Cterm), were non-functional in *let-653* mutants. N.S., not significant, p>0.01, two-tailed Fisher’s Exact test.

Next, we tested if the LET-653 C-terminus was sufficient for function or localization. Our attempts to make transgenic animals expressing only the LET-653 C-terminal tail failed, suggesting that the C-terminal tail is toxic (data not shown). Instead, to test C-terminus activity, we attached it to the ZPn domain (Figure 3C). This resulted in a secreted protein that did not rescue *let-653* mutant duct defects and did not localize to the duct lumen (Figure 3C, D). In the vulva, most LET-653(ZPn+Cterm) protein formed large, irregular clumps, but some of the protein localized along the vulva apical membrane (Figure 3C). These observations suggest that the C-terminus does have some capacity for aggregation and/or matrix binding. However, the full ZPc domain is required for tube shaping and appropriate apical matrix localization.

### LET-653 cleavage promotes ZPc function

Because ZP cleavage is thought to promote ZPn polymerization, we examined how it impacts LET-653(ZP) function. To test the role of cleavage in LET-653, we mutated the best candidate furin cleavage site (CCS) from RRYR to AAYA in a transgene encoding the full ZP domain (Figure 4A, B). Western blotting confirmed that, unlike wild-type LET-653, the mutated LET-653(ZP, AAYA) protein is not cleaved in *C. elegans* embryos (Figure 4F). Although this uncleavable protein still localized to the duct lumen, it localized weakly, or not at all, to the apical membrane in the vulva lumen (Figure 4B-D), and was unable to rescue *let-653* mutant lethality (Figure 4E). LET-653(ZP, AYAA) was expressed at similar levels and cleared from the duct lumen at the same time as the wild-type ZP domain (Figure 4C, Figure S2), indicating that differences in stability or clearance do not account for these results. We conclude that cleavage promotes LET-653(ZP) function and localization.

**Figure 4.**
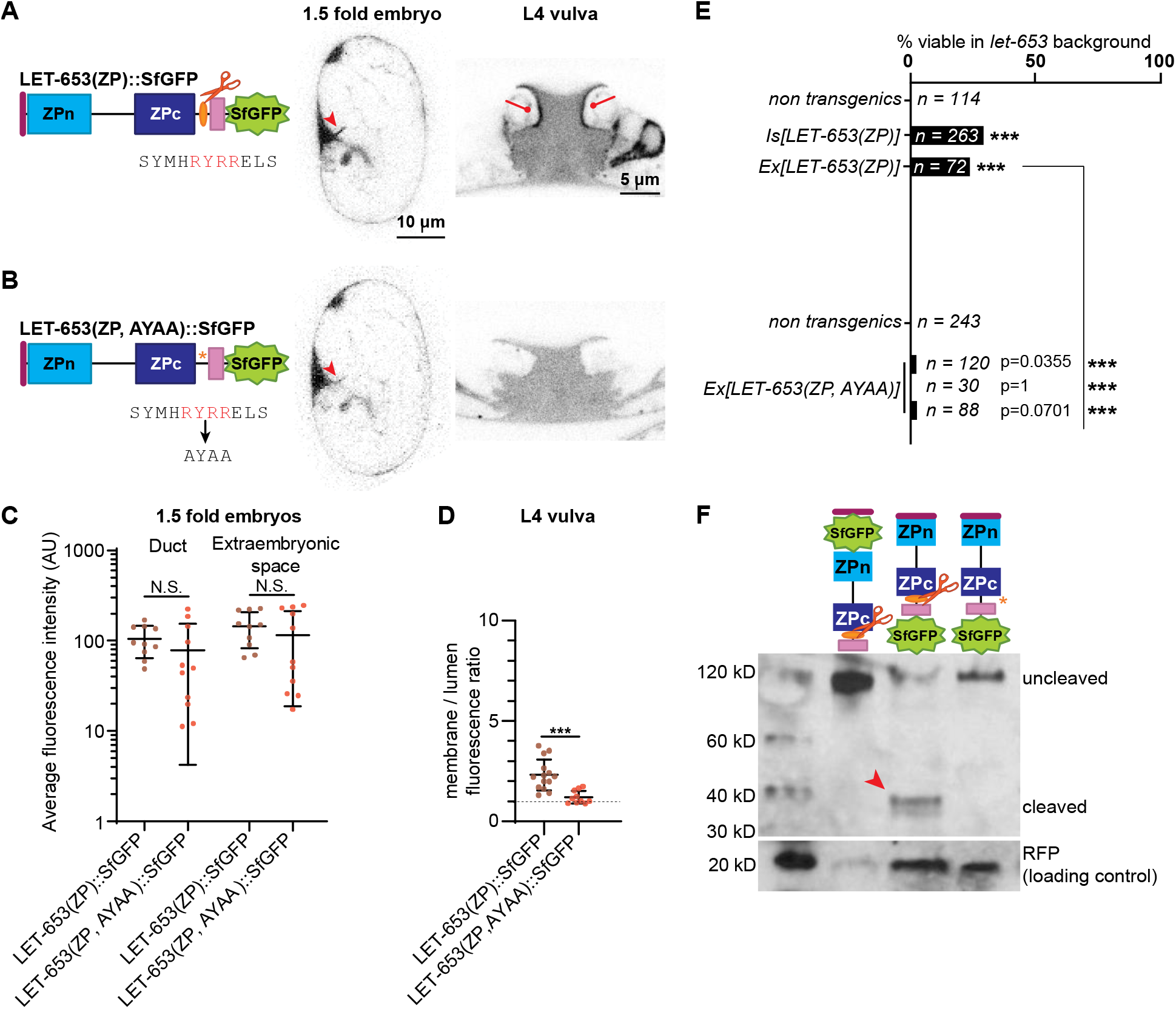
Cleavage promotes function and localization of the LET-653 ZP domain. A) LET-653(ZP)::SfGFP localized consistently to the embryonic duct lumen and the larval vulva dorsal apical membranes. B) LET-653(ZP, AYAA) cleavage mutant. The conserved CCS (red) was mutated to AYAA. Although LET-653(ZP, AYAA) was sometimes found in the embryonic duct lumen, it was never observed at vulva apical membranes. C) Duct and extraembryonic fluorescence in 1.5 fold embryos (see Figure 1). p>0.01, Mann Whitney two-tailed U test. Error bars; standard error. D) Ratio of apical membrane-associated to lumen fluorescence in the L4 vulva (see Figure 1). ***p<0.0001, Mann Whitney two-tailed U test. Dashed line indicates a ratio of 1, where membrane fluorescence is equal to luminal fluorescence. Error bars; standard error. E) LET-653(ZP)::SfGFP was partly functional in a *let-653* mutant background, though rescue was not as strong as that seen with N-terminally tagged transgenes (compare to Figure 2F). Three independent lines of LET-653(ZP, AYAA)::SfGFP had little activity. Although the comparisons to non-transgenic siblings are not statistically significant, the few survivors likely represent some biological function for LET-653(ZP, AYAA), as *let-653* mutants never survive (Gill et al., 2016). ***p<0.001, two-tailed Fisher’s Exact test. F) Western blot demonstrating abrogation of cleavage. SfGFP::LET-653(ZP) runs at ~120 kD while LET-653(ZP)::SfGFP runs at ~40 kD, indicating that LET-653 is cleaved at its C-terminus as previously reported (Gill et al., 2016). LET-653(ZP, AYAA)::SfGFP runs at nearly the same size as N-terminally tagged LET-653, indicating it is not cleaved. Loading control is RFP, which is co-expressed from transgenes containing LET-653 and whose expression is proportional to the amount of LET-653 in transgenic *C. elegans.* Blot is representative of n = 4 biological replicates.

To determine how cleavage enables LET-653(ZP) function, we mutated the cleavage site in a transgene expressing LET-653(ZPc). If, as previously proposed (Bokhove et al., 2016), cleavage relieves an inhibitory ZPn-ZPc interaction, then cleavage should no longer be required when the ZPn is absent. In contrast, if cleavage changes ZPc conformation to enable ZPc interactions, then cleavage should still be required in the absence of ZPn. LET-653(ZPc, AYAA) localized efficiently to the apical membrane of the vulva and to the duct lumen of embryos, and partially rescued *let-653* mutant duct defects (Figure 5B, C-E). Western blots confirmed that LET-653(ZPc, AYAA) was rarely cleaved (Figure 5F). We conclude that cleavage is not required for localization of the isolated ZPc domain but does promote its function. These results suggest that LET-653(ZPc) can bind to a relevant matrix partner without being cleaved, but that it requires cleavage for full or efficient function. According to this model, cleavage at the CCS relieves ZPn-dependent inhibition to facilitate binding to the first partner, but also changes other functional properties of the LET-653(ZPc) module.

**Figure 5.**
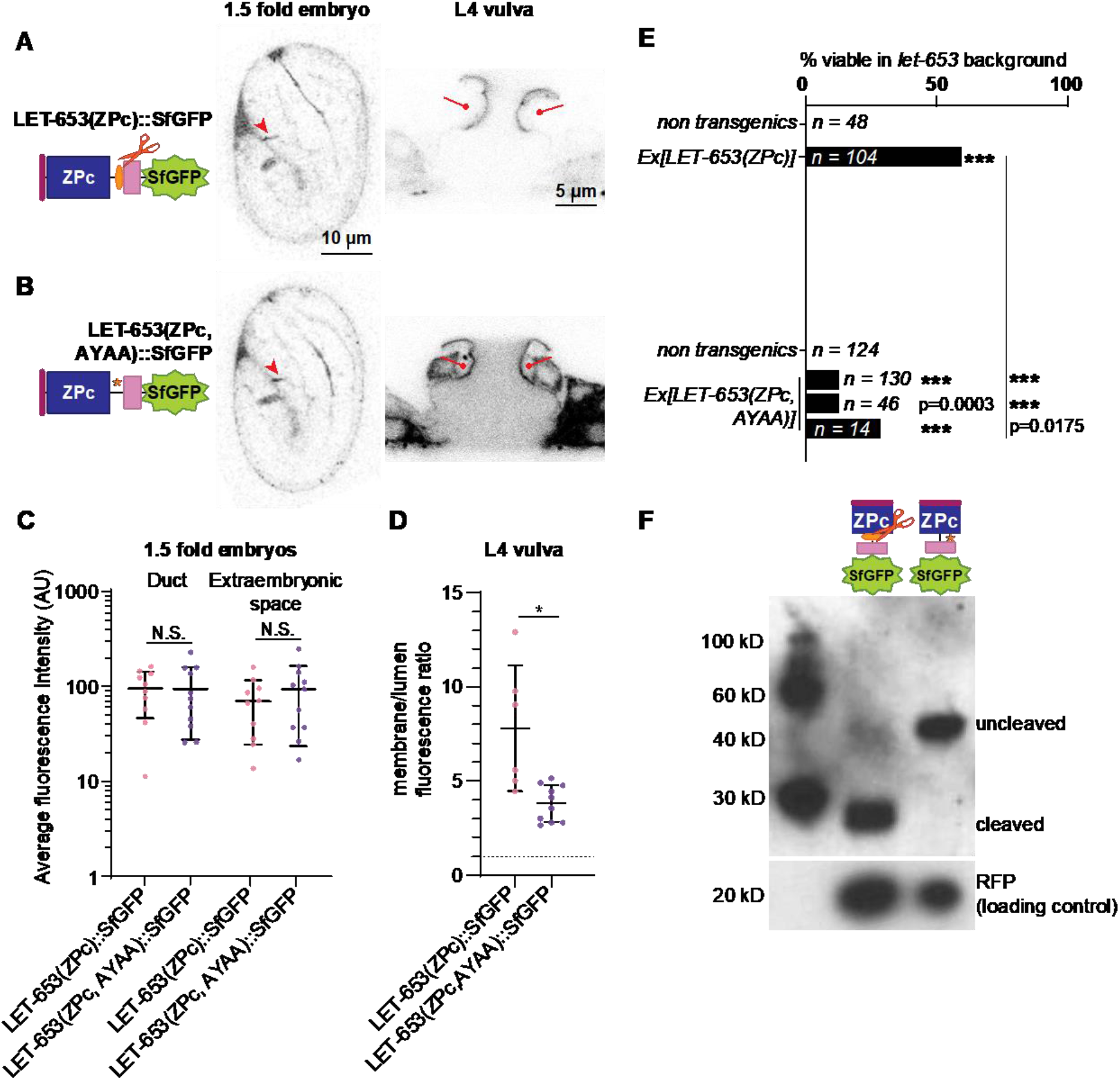
LET-653 cleavage relieves ZPn-dependent inhibition of ZPc. A) LET-653(ZPc)::SfGFP and B) LET-653(ZPc, AYAA)::SfGFP both localized to the duct lumen and vulva apical membranes. C) Duct and extraembryonic fluorescence in 1.5 fold embryos within boxes drawn on relevant structures (see Figure S1). p>0.01, Mann Whitney two-tailed U test. Error bars; standard error. D) Ratio of membrane-associated to lumen fluorescence in the L4 vulva (see Figure S1). *p=0.0047, Mann Whitney two-tailed U test. Dashed line indicates a ratio of 1, where membrane fluorescence is equal to luminal fluorescence. Error bars; standard error. E) LET-653(ZPc)::SfGFP rescued *let-653* mutants. Despite localizing normally, 3/3 lines of LET-653(ZPc, AYAA)::SfGFP only partially rescued *let-653* mutants. ***p<0.001, two-tailed Fisher’s Exact test. F) Western blot demonstrating abrogation of cleavage. LET-653(ZPc)::SfGFP runs at ~40 kD. Mutating the cleavage site (see Figure 4A) resulted in a protein that ran at ~60kD, close to the estimated size of the entire protein fragment based on amino acid sequence. Loading control is RFP (see Figure 4). Blot is representative of n = 3 biological replicates.

### LET-653(ZPc) incorporates stably into the aECM

To test if LET-653(ZPc) incorporates stably into the aECM, we performed FRAP analysis. In line with previous results (Gill et al., 2016), roughly 50% of LET-653(ZP) fluorescence was bleached and this fluorescence recovered poorly, indicating it is immobile in its aECM layer (Figure 6A). In contrast, only about 25% of LET-653(ZPc) fluorescence could be effectively bleached, though similar to LET-653(ZP), the bleached portion recovered poorly (Figure 6B-D). Resistance to bleaching could indicate that 1) A portion of LET-653(ZPc) is highly mobile and recovers before bleaching can be detected; or 2) LET-653(ZPc) integrates into an aECM structure that shields the fluorescent protein from bleaching. Notably, however, bleaching and recovery of LET-653(ZPc) were poor compared to what we previously reported for secreted SfGFP alone (Gill et al., 2016). In any case, poor recovery of the bleached portion of LET-653(ZPc) is consistent with a portion of the protein stably incorporating into an aECM layer.

**Figure 6.**
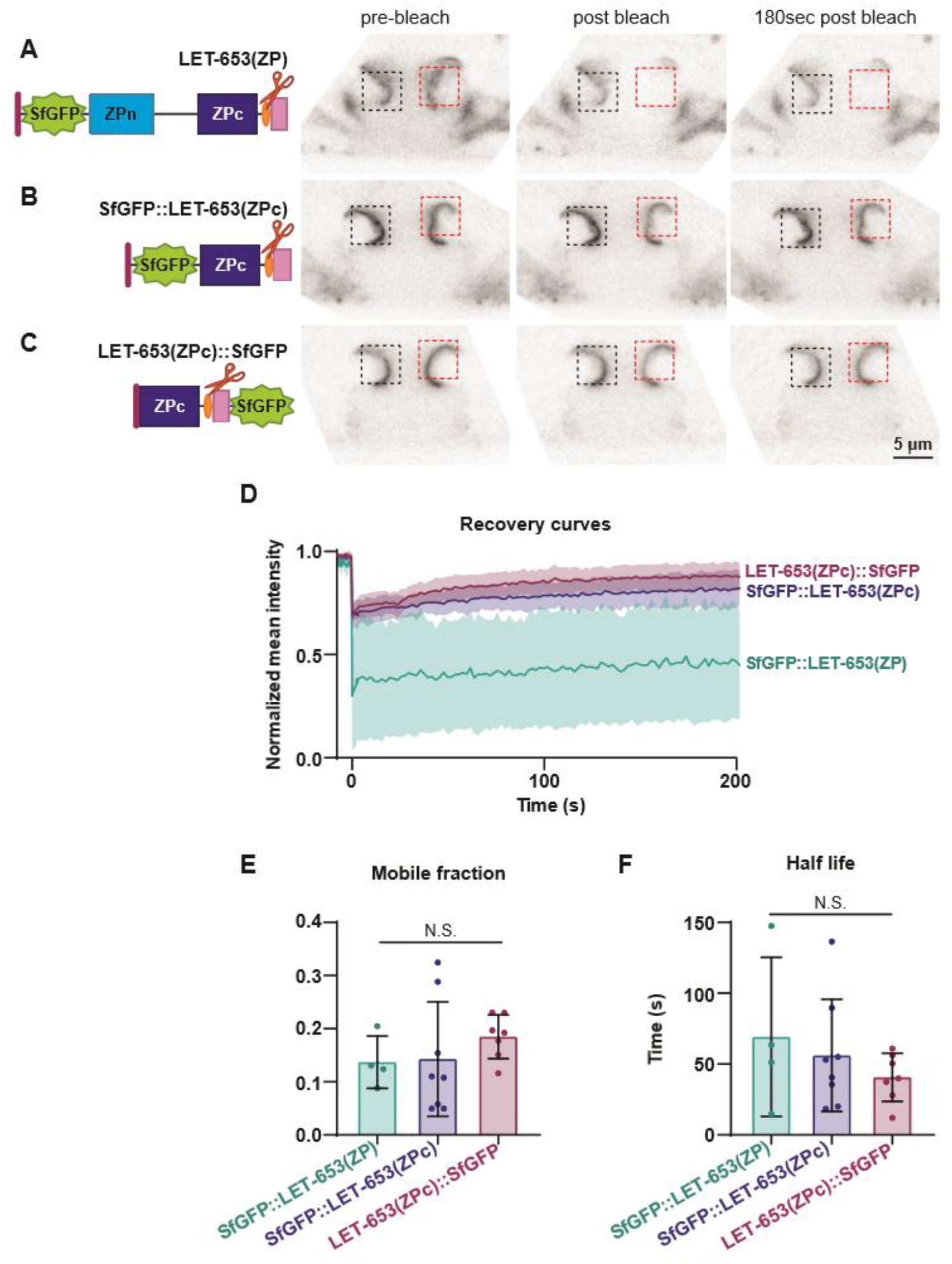
A portion of LET-653(ZPc) is immobile in the vulva aECM. Fluorescence recovery after photobleaching (FRAP) of LET-653 translational fusions in the duct lumen. Pre-bleach, bleach, and post-bleach frames taken from FRAP experiment on mid-L4 vulvas. Red box; bleached region of interest. Black box; unbleached region of interest. A) FRAP of SfGFP::LET-653(ZP). LET-653(ZP) is bleached but then recovers poorly. Representative of n = 4 replicates. B) FRAP of SfGFP::LET-653(ZPc). Representative of n = 8 replicates. C) FRAP of LET-653(ZPc)::SfGFP. Representative of n = 7 replicates. LET-653(ZPc) was not efficiently bleached regardless of the location of the SfGFP tag. D) Fluorescence recovery curves with mean and standard error for each LET-653 fusion protein. t = 0s represents the first post-bleach frame. E) Comparison of mobile fractions. No significant difference was measured between any groups (two tailed Mann-Whitney U test), though poor bleach efficiency could reflect a pool of highly mobile LET-653(ZPc). Error bars; standard error. F) Comparison of half time to recovery. No significant difference was measured between any groups (two tailed Mann-Whitney U test). Error bars; standard error.

### The LET-653 ZPc domain contributes to aggregates *in vitro*

To test whether the LET-653 ZPc domain polymerizes or aggregates, we expressed tagged LET-653 fragments in *Drosophila* S2R+ cells. Although, as previously observed, another *C. elegans* ZP protein, DYF-7, formed large fibrils on the surface of S2R+ cells (Low et al., 2019), LET-653 fragments formed no such structures and instead appeared diffuse (Figure S3). LET-653(ZP) or LET-653(ZPc) were efficiently secreted and properly cleaved by S2R+ cells, as assessed by Western blotting (Figure S3). However, using higher magnification, we observed large crystalline aggregates throughout the cell media when the LET-653 ZPc domain was present (Figure 7). Aggregates were rare and not fluorescent in media from S2R+ cells transfected with empty vector (EV) or SfGFP (Figure 7A-B, H-I), suggesting that endogenous S2R+-derived factors have little propensity for aggregation. In contrast, aggregates were much more common and moderately fluorescent in media containing LET-653(ZP) (Figure 7C, F, H-I). Strikingly, crystalline aggregates were highly abundant and brightly fluorescent in media containing LET-653(ZPc) (Figure 7E, H-I). Media containing LET-653(ZP, AYAA) had crystalline aggregates that were similar in brightness to those of media containing LET-653(ZP), but those aggregates were much less abundant (Figure 7F-I). We conclude that the LET-653 ZPc domain can oligomerize to form ordered aggregates, either on its own or in combination with other components derived from S2R+ cells or media. Furthermore, efficient aggregate formation *in vitro* correlates well with efficient matrix incorporation and function *in vivo*.

**Figure 7.**
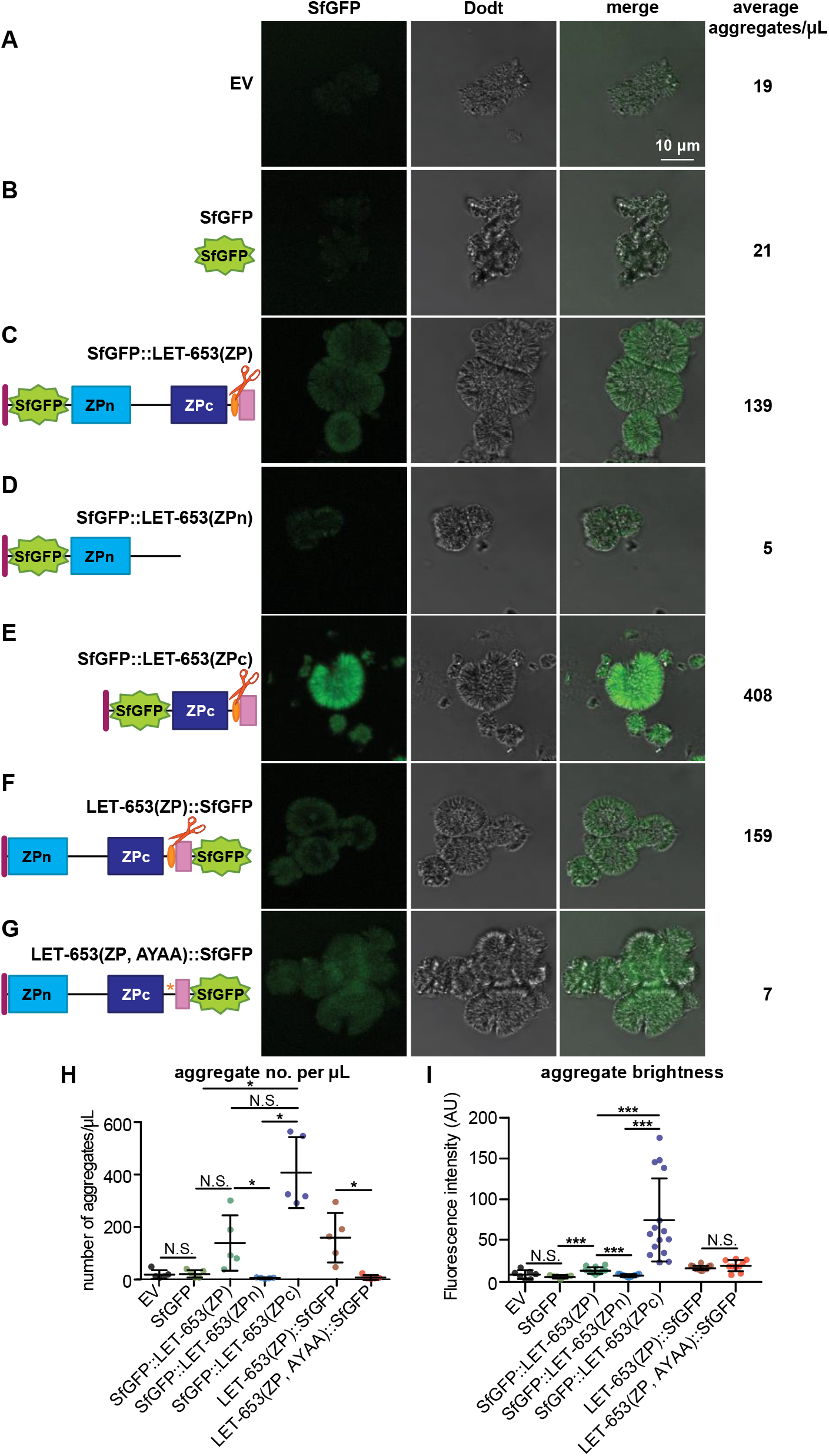
The LET-653 ZPc domain contributes to aggregates *in vitro*. A-G) Single confocal slices of freshly thawed cell media, with average aggregates per μl shown at right. A-B) Aggregates were rare and non-fluorescent in EV- or SfGFP-transfected cell media. C) Aggregates were more common and fluorescent in SfGFP::LET-653(ZP)-transfected media. D) SfGFP::LET-653(ZPn)-transfected media resembled negative controls; E) SfGFP::LET-653(ZPc) transfected cell media contained many aggregates of high fluorescence intensity. F-G) Although aggregates in media containing LET-653(ZP)::SfGFP and LET-653(ZP, AYAA)::SfGFP were similarly fluorescent, aggregates were much rarer in the latter sample. H) Quantification of aggregate density in media thawed and mounted on glass slides. *p<0.01, two-tailed Mann Whitney U test. SfGFP vs SfGFP::ZP, *p=0.0079. SfGFP vs SfGFP::ZPc, *p=0.0079. SfGFP::ZPc vs SfGFP::ZPn, *p=0.0079. SfGFP::ZP vs SfGFP::ZPn, *p=0.0079. ZP::SfGFP vs ZP, AYAA::SfGFP, *p=0.0079. Error bars; standard error. I) Quantification of aggregate fluorescence intensity. ***p<0.0001; two-tailed Mann-Whitney U test. Error bars; standard error. All experiments were performed on media from transfection 2.

To test the properties of the observed aggregates, we examined whether they stained with amyloid dyes or were heat sensitive. The dyes Nile Red and Congo Red stained aggregates whether secreted LET-653(ZPc) was present or not (Figure 8; and data not shown). These dyes bind amyloid structures, as well as hydrophobic materials, chitin, and some salt crystals (Greenspan, Mayer, & Fowler, 1985; Hawe, Sutter, & Jiskoot, 2008; Linder, 2018; Sackett & Wolff, 1987; Yakupova, Bobyleva, Vikhlyantsev, & Bobylev, 2019). To test heat sensitivity, we heated freshly thawed media to 50°C for fifteen minutes. Imaging revealed that aggregates remained after heating. However, brightly fluorescent aggregates were more numerous after cooling to 4°C for fifteen minutes (Figure 8C-D). The fact that these aggregates can be labeled with dyes and are non-heat-soluble indicates that they may have some biochemical characteristics of amyloid. Future studies are needed to determine the composition of these aggregates.

**Figure 8.**
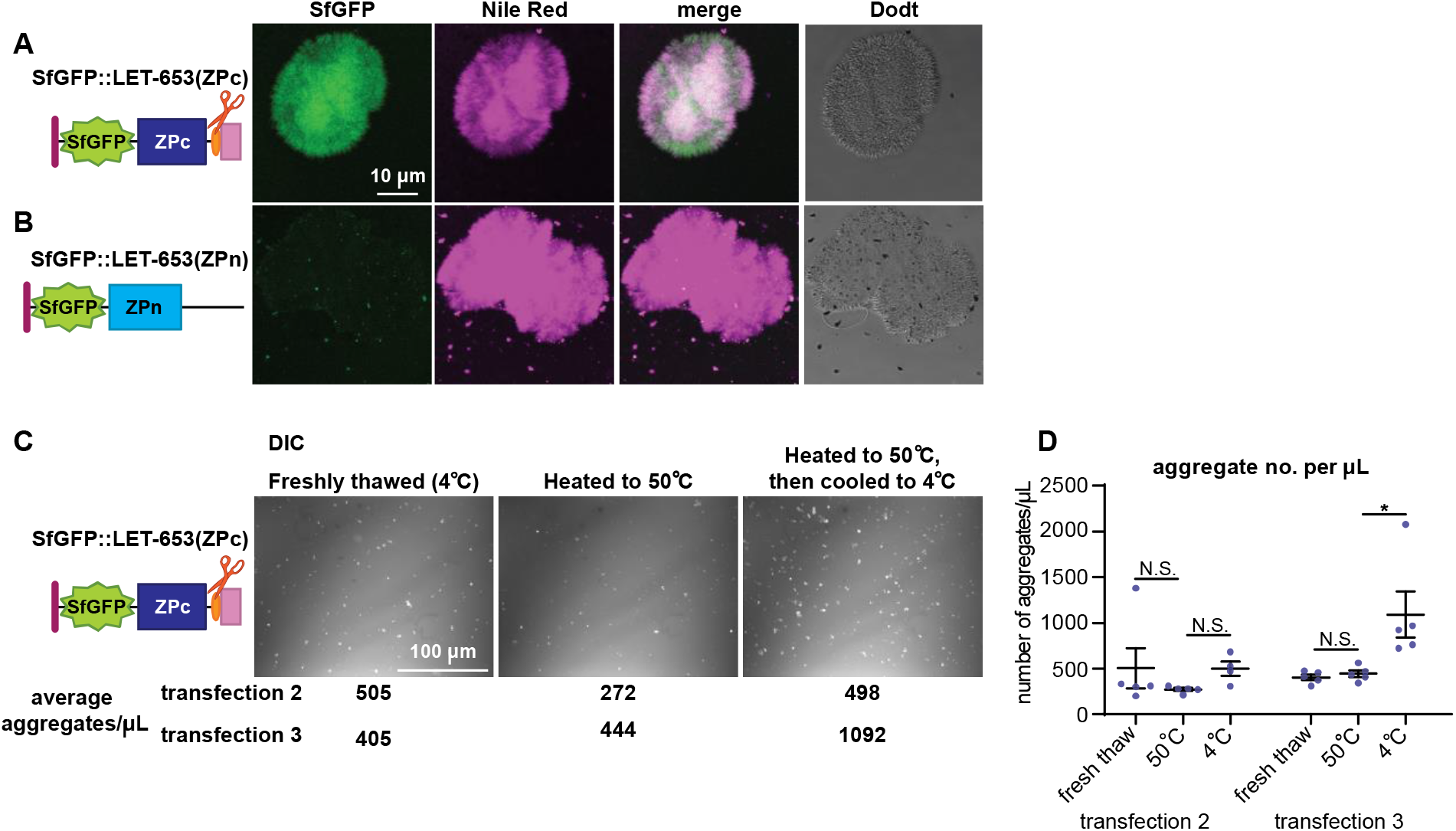
LET-653-bound aggregates stain with dyes and are not heat-labile. A-B) Aggregates stained with the lipid dye Nile Red regardless of whether LET-653(ZPc) was present. A) LET-653(ZPc) and Nile Red co-labelled aggregates. B) When LET-653(ZPc) was absent, aggregates, which were rare, still stained strongly with Nile Red. C) DIC images of LET-653(ZPc) aggregates. Aggregates were plentiful in freshly thawed media and remained after media was incubated for 15 minutes at 50 °C. Aggregate number increased somewhat after media was re-cooled to 4 °C for 15 minutes. D) Quantification of aggregate number. *p<0.01, two-tailed Mann Whitney U test. LET-653(ZPc) media heated to 50 °C vs LET-653(ZPc) media cooled to 4 °C, *p=0.0079. Error bars; standard error.

## Discussion

Although ZP proteins are critical components of aECMs in metazoans, how ZP proteins assemble in aECMs remains poorly understood. In this study, we investigated how one *C. elegans* ZP protein, LET-653, assembles into aECMs. We found that, counter to prevailing models for ZP assembly, LET-653 functions and localizes via its ZPc domain (Figure 9A-B). Cleavage of the C-terminus serves to release inhibition of the ZPc by the ZPn domain, but also likely promotes conformational changes in the ZPc domain that allow it to function efficiently. *In vitro*, LET-653(ZPc) spontaneously incorporates into large aggregates. These data demonstrate that ZPc domains can form higher order structures and assemble in aECMs, functions that were previously attributed only to ZPn domains.

**Figure 9.**
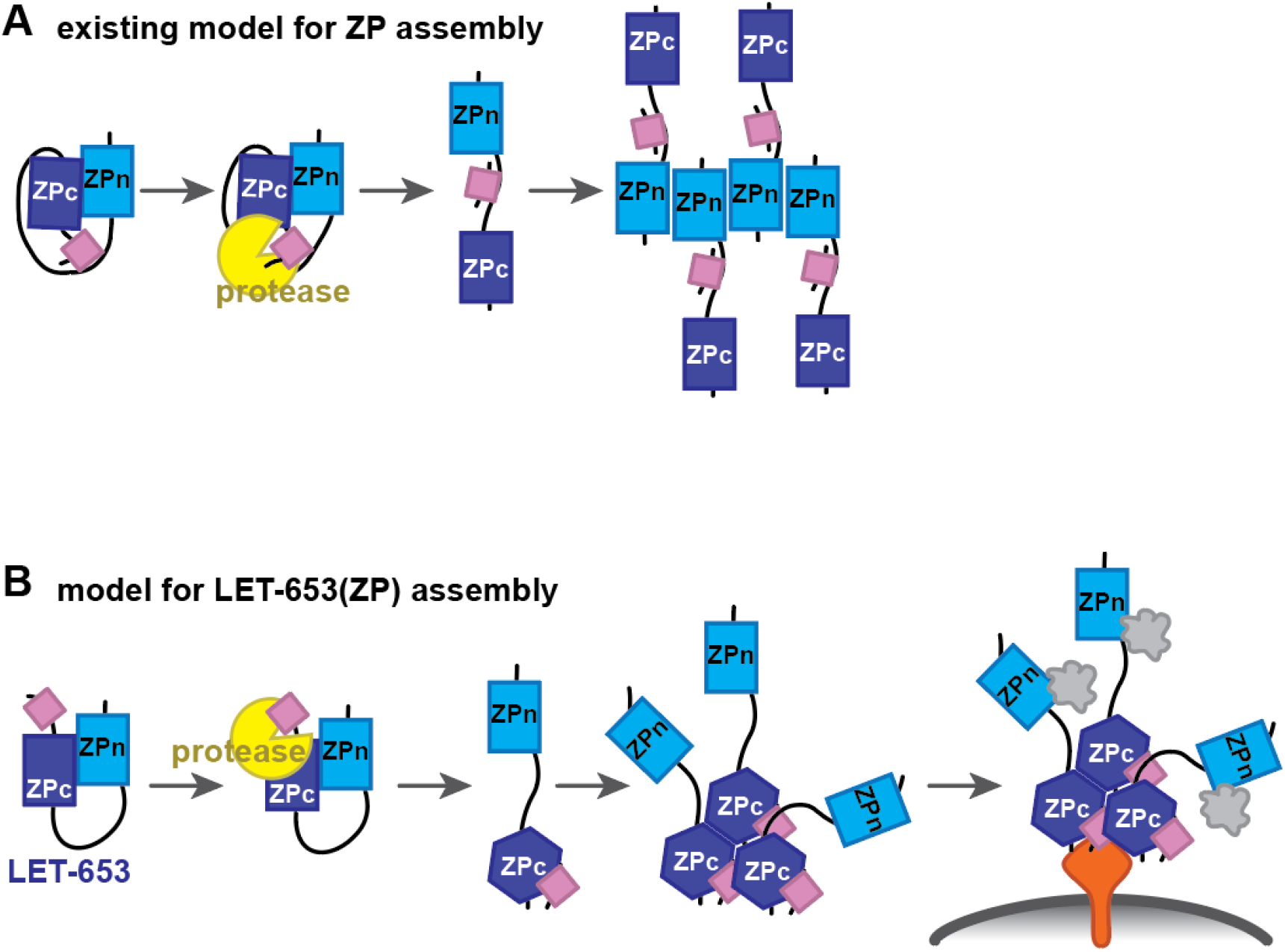
Model for LET-653 matrix assembly. A) Traditional model for ZP assembly. Prior to cleavage, the hydrophobic C-terminus interacts with a hydrophobic patch in the linker. This interaction causes the ZPc domain to inhibit the ZPn domain from interacting with itself or other partners. After the C-terminus is cleaved, the ZPn domain is free of inhibition and polymerizes. B) Model for LET-653 assembly. Prior to cleavage, the ZPn domain inhibits the ZPc domain. After cleavage, the C-terminus associates with the ZPc domain, which changes conformation. The ZPn domain no longer inhibits the ZPc domain. The ZPc domain oligomerizes and binds a membrane-anchored partner(s) (orange). The ZPn domain may bind a partner(s) (gray cloud) that prevents it from inhibiting ZPc aggregation. Membrane; gray line.

### ZPc domains can function independently of ZPn domains

ZPc domains have been generally assumed to support ZPn domains and not to function independently (Bokhove & Jovine, 2018). Initial surveys of ZP proteins across phyla suggested that some contain ZPn domains without ZPc domains, and yet are still capable of polymerization (Jovine et al., 2006), while proteins with only ZPc domains were not thought to exist (Bokhove & Jovine, 2018). However, we found that *C. elegans* does have five predicted ZPc-only proteins (Cohen et al., 2019). In proteins like uromodulin that have polymerizing ZPn domains, ZPc domains are thought to inhibit the ZPn domain (Bokhove & Jovine, 2018). In the TGFβ receptors Betaglycan and Endoglin, ZPc domains dimerize and can bind ligands, but have not been reported to form higher-order structures (Diestel et al., 2013; Lin et al., 2011; Saito et al., 2017). Our results indicate that ZPc domains can function independently of ZPn domains for both matrix incorporation and tissue-shaping.

In fact, in the case of LET-653, the ZPn domain inhibits ZPc activity, perhaps to confine ZPc activity to precise times and places within the aECM. We did not detect any independent role for the ZPn domain in shaping the duct lumen, and this domain is not able to localize specifically in the vulva aECM. This result is surprising given the centrality of ZPn domains to most models of ZP function (Bokhove & Jovine, 2018). The LET-653 ZPn domain lacks a key tyrosine residue present in most polymerizing ZP proteins (Gill et al., 2016; Monne et al., 2008), which could explain functional differences between the LET-653 ZPn domain and others.

Intriguingly, this tyrosine residue is also absent in a number of other *C. elegans* ZP proteins (Cohen et al., 2020), the *Drosophila* protein Zye (Fernandes et al., 2010) and some mammalian proteins including Endoglin (Monne et al., 2008) and Oit3 (Figure S4). Researching additional ZP proteins may reveal that both ZPn and ZPc domains can have diverse functions, and that either domain, or both domains simultaneously, can direct matrix incorporation.

The LET-653 ZPc domain has additional cysteines compared to some ZP proteins. ZPc domains fall into several families, with 4, 6, or 8 cysteines (Bokhove & Jovine, 2018; Cohen et al., 2019). The LET-653 ZPc domain has 8, as does the Zona Pellucida ZP protein ZP3 (Bokhove & Jovine, 2018; Han et al., 2010). Like LET-653, ZP3 has a cleaved C-terminus that can bind the ZPc domain (Han et al., 2010). It is possible that all ZPc domains with 8 cysteines have analogous assembly mechanisms. Testing the assembly of additional proteins in this class may expose patterns in ZP structure and function.

### The LET-653 ZPn domain inhibits two separable aspects of ZPc function

Intramolecular interactions between ZPn and ZPc domains during cellular trafficking are thought to hold ZP proteins in an inactive conformation to prevent premature oligomerization (Bokhove & Jovine, 2018). C-terminal cleavage, perhaps by proteases at the plasma membrane, relieves this auto-inhibition to allow matrix incorporation (Boja et al., 2003; Brunati et al., 2015; Jimenez-Movilla & Dean, 2011). Our data are consistent with this model, although the relative roles of the ZPn vs ZPc domains are reversed. Our data further suggest that there are two distinct roles of the ZPc domain that are subject to auto-inhibition: 1) the ability to bind to a matrix partner that determines spatial specificity; and 2) some other change (possibly oligomerization) that is essential for tube-shaping function.

The relevant matrix partner(s) of LET-653(ZPc) are not known, but they appear to be located on the apical surfaces of the excretory duct tube and vulE and vulF vulva cells. Although *let-653* is expressed by all vulva cells (Gill et al., 2016), genetic perturbations to increase or eliminate vulE or vulF cell types showed that these cells have a specific ability to recruit LET-653(ZP) (Cohen et al., 2020). Furthermore, vulF cells contain large matrix-secreting vesicles whose contents appear disorganized in *let-653* mutants, suggesting that LET-653 may traffic through the same compartment with its partner(s) (Cohen et al., 2020). Identifying the relevant partner(s) may require a combination of genetic and biochemical approaches and would likely offer crucial insight into how LET-653 incorporates into matrix and shapes tubes.

### The LET-653 ZPc domain may form oligomers

The second ZPc function enabled by cleavage might be oligomerization. A portion of LET-653(ZPc) appears immobile in the aECM *in vivo* (Figure 6) and incorporates into crystalline aggregates *in vitro* (Figure 7). Although the *in vitro* aggregates could be multicomponent structures containing other S2R+ cell-derived proteins and/or salts, their formation, growth or stability is enhanced by LET-653(ZPc) (Figure 7) and their number correlates well with *in vivo* matrix incorporation and function. LET-653 proteins that are not functional *in vivo*, including uncleavable LET-653(ZP, AYAA), did not increase the number of aggregates (Figure 7). Cleavage is required for efficient function of the isolated ZPc domain, even though it is not required for its proper localization (Figure 5). Together, these data suggest that LET-653(ZPc) may oligomerize to form its matrix layer *in vivo,* and that cleavage not only relieves ZPn-dependent inhibition of this activity, but also promotes oligomerization more directly.

Our data indicate that the cleaved C-terminus stays associated with the ZPc domain, since LET-653(ZPc) tagged at either end results in identical localization patterns. Models for ZP organization generally show the C-terminus binding to the linker (Bokhove et al., 2016), but the crystal structure of the protein ZP3 indicates the C-terminus can bind the ZPc domain (Monne et al., 2008). Furthermore, the C-terminus alone can confer aggregation properties when attached to the ZPn domain (Figure 3). We propose that a cleavage-dependent change in the relative conformations of the main ZPc and C-terminus may facilitate oligomerization and interactions with other partners (Figure 9B).

*In vitro*, LET-653(ZP) incorporates less efficiently into aggregates than LET-653(ZPc) despite efficient cleavage, demonstrating that, in some contexts, the ZPn domain could continue to inhibit the ZPc domain after cleavage. *In vivo,* LET-653(ZPn) may have a binding partner that prevents it from associating with the ZPc after cleavage (Figure 9B). This partner could be another protein or lipid or could be the N-terminal LET-653 PAN domains (Figure 1A), which were not included in these experiments, but which normally limit ZP-dependent matrix incorporation (Gill et al., 2016). In summary, we propose a model in which ZPn domain inhibition of the ZPc domain is relieved by cleavage and the ZPn domain is maintained away from the ZPc domain by as-yet-unknown binding partners of each domain (Figure 9B).

LET-653 incorporates into roughly spherical aggregates that resemble salt crystals. These aggregates may consist largely of protein or could represent protein-salt complexes. Intriguingly, salt appears to stimulate uromodulin to form filaments (Bokhove et al., 2016). Furthermore, the ZP protein Oit3 (also known as LZP), a proposed partner of uromodulin, can form spherical aggregates of unknown organization *in vitro* (Shen et al., 2009). However, how these Oit3 aggregates assemble, if they include other co-factors, such as uromodulin or salts, and whether they form *in vivo* is not clear. Previous *in vitro* studies of ZP proteins have been challenging, as many cell types fail to cleave or secrete intact ZP proteins appropriately (Bokhove et al., 2016; Han et al., 2010). S2R+ cells fortuitously express a compatible protease and offer a straightforward model to assess ZP assembly and to characterize ZP aggregates.

### Conclusion

ZP proteins are a diverse family of aECM components that are found across the animal kingdom. ZP proteins shape many apical surfaces (Plaza et al., 2010), changes in ZP protein expression or structure are implicated in the evolution of morphological features (Smith, Davidson, & Rebeiz, 2020) and speciation (Raj et al., 2017), and ZP protein loss contributes to multiple human diseases. This work harnessed *C. elegans* to demonstrate a new mode of ZP protein assembly in which the ZPc domain is the primary functional unit. Future work should take advantage of additional tractable models to characterize the true diversity of ZP protein function and assembly mechanisms.

## Materials and Methods

### Worm strains, alleles and transgenes

All strains were derived from Bristol N2 and were grown at 20°C under standard conditions (Brenner, 1974). *let-653* mutants were obtained from mothers rescued with a *let-653*(+) transgene (Gill et al., 2016). Mutants used were *let-653(cs178)* (an early nonsense allele) (Gill et al., 2016). Strain generated using CRISPR/Cas9 was *let-653(cs262[LET-653::SfGFP])* (Cohen et al., 2020). See Table S1 for a complete list of strains generated in this study.

### Staging and microscopy

Embryos were selected at the 1.5-fold stage based on morphology. Larvae were staged by size and vulva lumen morphology. Fluorescent, brightfield, Differential Interference Contrast (DIC), and Dodt (an imaging technique that simulates DIC) (Dodt & Zieglgansberger, 1990) images were captured on a Zeiss Axioskop compound microscope fitted with a Leica DFC360 FX camera or with a Leica TCS SP8 confocal microscope (Leica, Wetzlar Germany). Images were processed and merged using ImageJ.

### Quantification of LET-653 fluorescence

To confirm that LET-653 transgenes were adequately expressed, their fluorescence intensity was measured in embryos within boxes drawn on single confocal slices. Boxes were drawn either to fit the small duct lumen using the ImageJ “Rotated rectangle” tool or were drawn to roughly 3×3 microns within the extraembryonic space (Figure 1D). The average fluorescence intensity per pixel within those regions was measured using the ImageJ “Mean gray value” tool.

Measurements of LET-653 enrichment at the vulva vulE and vulF cell apical membranes were performed as in Bidaud-Meynard et al. (Bidaud-Meynard, Nicolle, Heck, Le Cunff, & Michaux, 2019). 10 pixel-thick-lines were drawn across the dorsal region of the vulva, including both apical membranes. 5 pixel values representing LET-653 fluorescence intensity at each apical membrane (totaling 10 values) were averaged and divided by 10 pixel values from the luminal region directly between the vulE and vulF apical membranes (Figure 1G). All measurements are taken from single confocal Z-slices. Final values reported are ratios.

### *let-653* transgenes

All *let-653* transgenes were generated by inserting LET-653 fragments into the plasmid pJAF5, which contains a 2.2 kb fragment containing the *let-653* promoter and upstream regulatory region (Gill et al., 2016). All plasmids were generated via restriction enzyme cloning and were verified by Sanger sequencing. *let-653* encodes three isoforms of a protein with two N-terminal PAN/Apple domains and a ZP domain. As the shortest isoform, LET-653b, was previously shown to be sufficient for function, all structure-function constructs are based on LET-653b (GenBank accession no: X91045.1) (Gill et al., 2016). Plasmids were co-injected at concentration 17 ng/μL with marker pHS4 (*lin-48pro*::mRFP) at concentration 50 ng/μL. Integrated transgenes were generated using TMP/UV as described (Nance & Frøkjær-Jensen, 2019) and include: *csIs92 [let-653pro::SfGFP::LET-653(ZPc); lin-48pro::mRFP], csIs95 [let-653pro::LET-653(ZP, AYAA)::SfGFP; lin-48pro::mRFP], csIs96 [let-653pro::LET-653(ZP)::SfGFP; lin-48pro::mRFP].* In most cases, two or three independent lines of each transgene were assayed for rescue activity and gave similar results. Only a single line per transgene was used for the image quantifications shown. A complete list of transgenes, primers, and plasmids generated for this study can be found in Table S2.

### FRAP

Specimens were mounted on 10% agarose pads containing 20 mM sodium azide and 10 mM levamisole in M9 after Gill et al., 2016 (Gill et al., 2016). FRAP was performed using Leica Application Suite X software FRAP module on a Leica TCS SP8 MP confocal microscope. Three regions of interest of 3 × 3 μm were defined within the wizard: a bleached region of interest (bleached ROI), an unbleached ROI of similar fluorescence intensity, and a background ROI drawn in a region that did not include an animal. Maximum laser intensity was used for bleaching experiments and experiments were carried out over the course of several weeks. Mean fluorescence intensities within the ROIs were measured at specified intervals. Background ROI values were subtracted from both the bleached and unbleached ROI values. Corrected bleach ROI values then were divided by corrected unbleached ROI values (bleached - background / unbleached - background), and the resulting ratios were normalized such that the maximum intensity was set to 1. The following experimental time-course was used: 20 prebleach frames every 0.4 sec, 10 bleach frames every 0.4 sec, and 90 postbleach frames every 2.0 sec. Pre- and postbleach laser intensity was set to 1% and bleach laser intensity was set to 100%. FRAP plots were created and analyzed using Prism (Graphpad, San Diego California), where one-phase association curves derived from the model Y = Y0 + (Plateau-Y0)*[1-e^(−Kx)] were fitted to the data. For statistical tests, mobile fractions and recovery half-times were derived from one-phase association curves fitted to individual experiments. Mobile fraction = Plateau-Y0; t1/2 = ln(2)/K, where K is the recovery rate constant.

### Drosophila cell culture and media experiments

S2R+ cells were cultured under standard conditions (Yanagawa, Lee, & Ishimoto, 1998). Cells were transfected using Effectene (Qiagen, 301427, Venlo Netherlands). Plasmids for transfection were generated by inserting cDNAs containing *let-653b* into pAC5.1 as XbaI - EcoRI fragments alongside the *unc-54* 3’ UTR as an EcoRI - ApaI fragment. Three sets of S2R+ cells were independently transfected with 1 μg DNA of the indicated constructs. Cells were grown under standard conditions for 72 hours before imaging and media collection. Cells were imaged under brightfield and fluorescence microscopy using a Leica SP5 confocal microscope, and media and cells were then collected for further analysis. Media was collected by pipetting and was then centrifuged for 1 min at 10,000. Cells were washed in PBS then lysed via incubation for 20 minutes at 4 °C in 1 ml lysis buffer (50 mM Tris-HCl pH 7.4, 500 mM NaCl, 1% NP40, 1 mM EDTA, 0.25% sodium deoxycholate, 1 mM sodium orthovanadate, 1× complete protease inhibitor cocktail (Roche, Basel Switzerland) and 1 mM PMSF). Cell lysates were centrifuged for 20 minutes at 4 °C at 10,000 rpm to remove debris. Media and cell lysates were stored at −80 °C until ready for use. All three sets of transfections showed evidence of LET-653 secretion and cleavage as assessed by Western blotting. Two out of two tested sets of transfections showed evidence of ZPc aggregation.

To image aggregates, 3 μL freshly thawed media was pipetted onto glass slides. DIC images for counting aggregates were obtained with a Zeiss Axioskop. 5 evenly spaced 92 mm x 92 mm fields of view were imaged per slide. To quantify aggregate number, DIC images were first subjected to a grayscale threshold filter in ImageJ (default threshold; limits: 85-255) and then aggregates were counted using the 3D Particle Counter plugin. The number of aggregates per field of view was adjusted to represent the number of aggregates per μL. To measure fluorescence intensity of aggregates, a box of 5 x 5 microns was drawn within aggregates and the average intensity per pixel (Mean gray value) was recorded from a single confocal Z-slice. A minimum of 5 aggregates per sample were measured.

To measure aggregate stability at different temperatures, 12 μL of media was thawed and aliquoted. 3 μL of this media was immediately pipetted onto glass slides for imaging. The remaining 12 μL of media was heated to 50°C in a thermocycler for 15 minutes and then a portion was immediately imaged. Another portion was reserved and cooled to 4°C in a thermocycler for 15 minutes before imaging. Aggregate number from DIC images was quantified as described above, except that before thresholding, images were processed using the ImageJ plugin “Background Correction”. To stain aggregates with Nile Red or Congo Red, a solution of 100 mM in acetone or water, respectively, was mixed with media for a final concentration of 1 mM and the sample was then imaged immediately.

Western blotting was performed on 5 μL of media or cell lysate as described below.

### Western blotting

Early embryos were collected via the alkaline bleach method (Stiernagle, 2006) from two 60 mm plates of nearly confluent worms and embryos were allowed to develop in M9 for 4-5 hours until they reached 1.5-fold stage. Worms were pelleted and transferred to Laemmli buffer (Bio-Rad, 161-0737) with 1x Protease Inhibitor Cocktail (Sigma, 2714) and 1:20 β-mercaptoethanol. Samples were transferred to −80C until ready for use. Samples were boiled for 10 minutes and then loaded into Mini-Protean TGX gradient gel (Bio-Rad, 456-1084) or a 15% acrylamide gel. Electrophoresis was performed at 0.02-0.03A under 1x electrophoresis buffer (Bio-Rad, 161-0732). Protein was transferred onto 0.2um nitrocellulose membrane (Bio-Rad, 162-0147) overnight in 1x transfer buffer (20% ethanol, 0.58% Tris Base, 2.9% Glycine, 0.01% SDS). Membranes were washed in PBS + 0.02% Triton (PBST), blocked for 1 hour at room temperature in PBST + 10% dry, nonfat milk, and then probed for GFP for 1 hour at room temperature with PBST + 10% milk + 1:1000 anti-GFP antibody (Rockland Immunochemicals, 600-101-215). To visualize the signal, Membranes were washed in PBST, probed with PBST + 10% milk + 1:5000 anti-Goat-HRP antibody (Rockland Immunochemicals, 605-4302) for 1 hour at room temperature and then washed with PBST before detection using SuperSignal West Femto Maximum Sensitivity Substrate (Pierce 34095) and film. Loading controls were carried out as described above using 1:1000 anti-RFP antibody (Abcam, ab62341) and 1:10,000 anti-rabbit-HRP antibody (GE Healthcare, NA934V). RFP corresponds to the *lin-48p::mRFP* co-injection marker present in all transgenes, and therefore reflects the number of transgenic animals present in each sample.

## Acknowledgements

We thank Andrea Stout (UPenn CDB Microscopy Core) for training and assistance with confocal imaging and interpreting FRAP data, Kelly Sullivan and Greg Bashaw for assistance with S2R+ cell experiments, Luca Jovine for generous advice and encouragement, and Erfei Bi and David Raizen for helpful discussions and comments on the manuscript. Some strains were provided by the CGC, which is funded by the NIH Office of Research Infrastructure Programs (P40 OD010440). This work was funded by NIH grants R01GM58540 and R01GM125959 to M.V.S.,T32 GM008216 and T32 AR007465 to J.D.C., R01 EB028320 and R35 GM128748 to M.C.B. and NSF iSuperseed grant 1720530 to M.C.B.

## Author Contributions

JDC conceived of and performed experiments, arranged figures, and wrote the paper. JGB performed experiments.

MCG offered critical advice and facilities, conceived of experiments, and interpreted data. MVS conceived of experiments, interpreted data, arranged figures and wrote the paper.

## Supplemental materials

**Figure S1.**
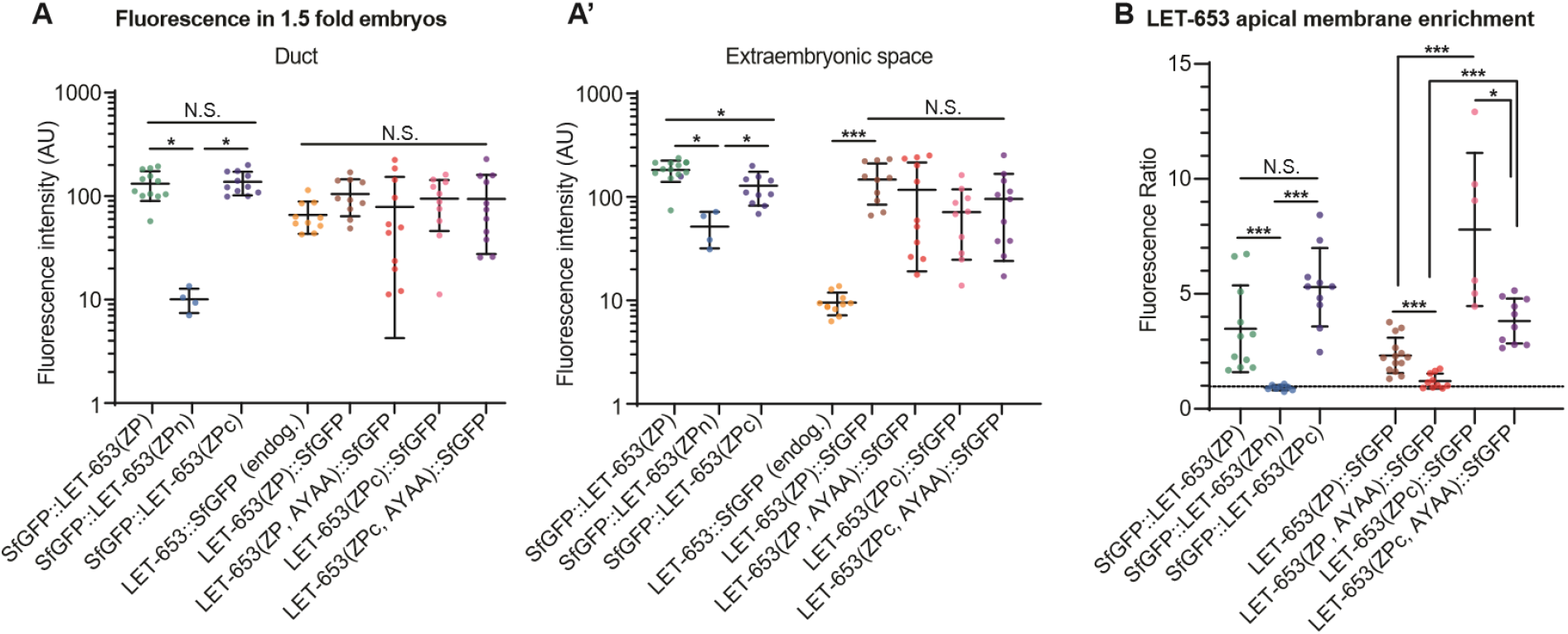
Method of measuring LET-653 in embryos and vulva. A-A’) Levels of LET-653 fluorescence in the 1.5 fold embryo. Functional transgenes were generally expressed at similar levels to endogenous LET-653 in the duct lumen, but accumulated to much higher levels in the extraembryonic space. Error bars; standard error. B) Quantification of membrane-associated vs. luminal LET-653 in the mid-L4 vulva for all groups. C-terminally and N-terminally tagged LET-653(ZP) and LET-653(ZPc) are not significantly different from one another. However, LET-653(ZPc, AYAA) is significantly more enriched at the apical membrane than LET-653(ZP, AYAA), and this result correlates with increased function in LET-653(ZPc, AYAA) vs LET-653(ZP, AYAA) (Figures 4-5). Dashed line indicates a ratio of 1, where membrane fluorescence is equal to luminal fluorescence. Error bars; standard error.

**Figure S2.**
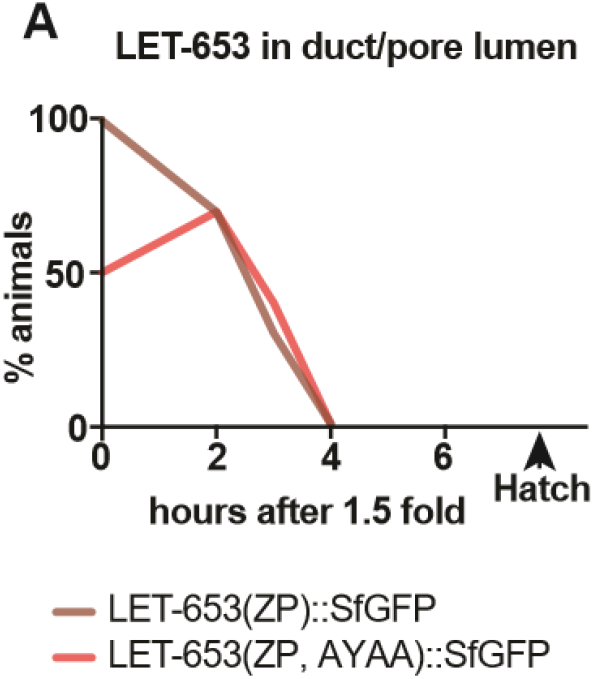
LET-653(ZP, AYAA) is cleared normally from the embryonic duct. A) LET-653(ZP) and LET-653(ZP, AYAA) are both present in the duct lumen at 1.5 fold stage but are gradually cleared over several hours. Presence of LET-653 in the duct lumen was assessed visually via epifluorescence microscopy. LET-653(ZP, AYAA) was sometimes too faint to be detected within the duct lumen at the 1.5 fold stage. n = 20 for each time point.

**Figure S3.**
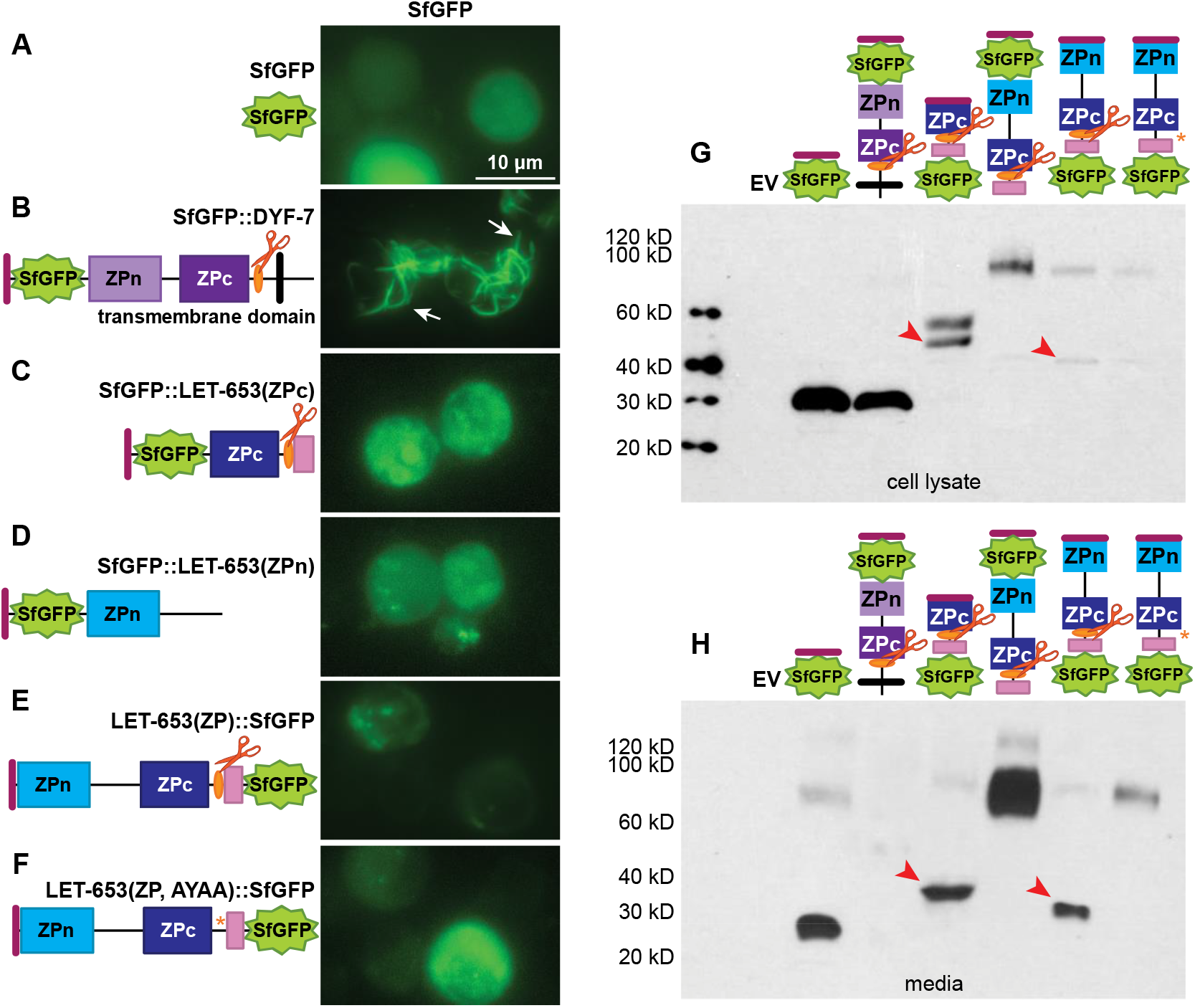
LET-653 is cleaved when expressed from S2R+ cells. S2R+ cells were transfected with 1 μg DNA of the indicated constructs. A-F) Epifluorescent images of cells 72 hrs post-transfection, representative of >100 cells per condition. Large intracellular or cell-bound fibrils (arrows) were visible from SfGFP::DYF-7 (Low et al., 2019) (B), but not from SfGFP alone or any LET-653 protein fusion. Black bar; transmembrane domain. Purple boxes; DYF-7 ZP domain. G) Western blot of S2R+ cell lysate probed with anti-GFP. Both un-cleaved and cleaved versions (arrowheads) of LET-653(ZP) and LET-653(ZPc) were visible. H) Western blot of S2R+ cell media probed with anti-GFP. LET-653(ZP) and LET-653(ZPc) were efficiently cleaved and cleavage products were visible in the media (arrowheads). LET-653(ZP, AYAA) was not cleaved but was still secreted into the media. EV; Empty vector. Western blot of media is representative of three blots from the three independent transfection experiments. Western blot of cell lysates was performed using cells from transfection 2. All images shown are of cells or media from transfection 2.

**Figure S4.**
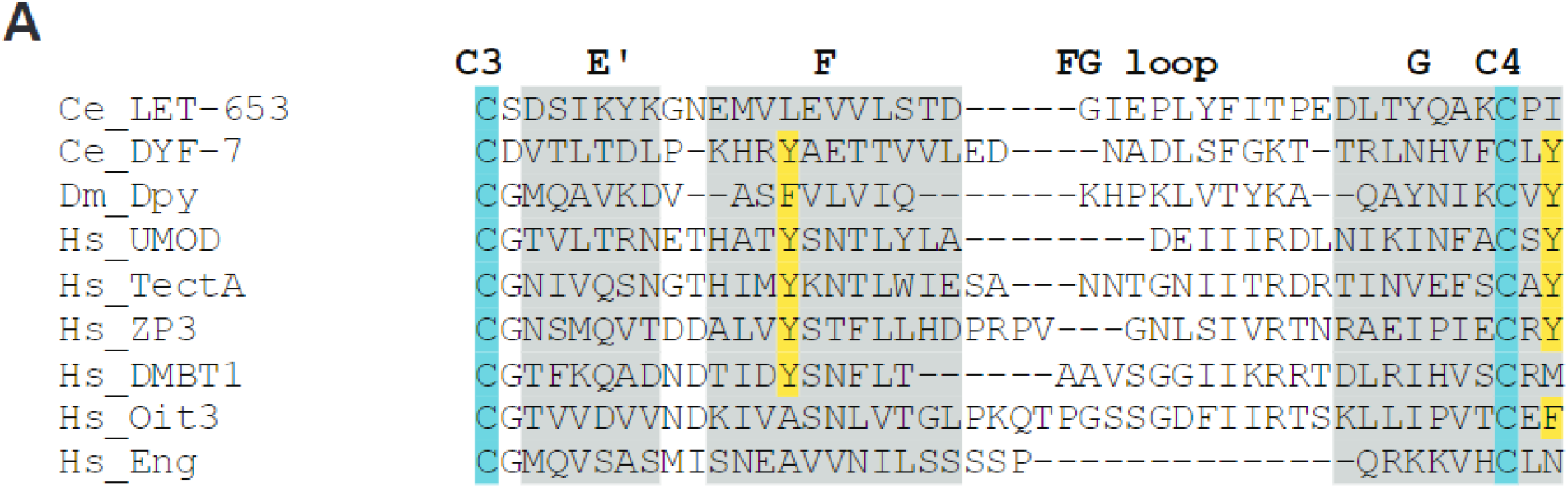
The human ZP proteins Oit3 and DMBT1 lack key tyrosine residues. Alignment of ZPn domains based on Monne et al., 2008 (Monne et al., 2008) and Gill et al., 2016 (Gill et al., 2016). Grey shading corresponds to known beta-strands in ZP3, blue indicates conserved cysteines and yellow indicates conserved aromatic residues. Like LET-653, Oit3 lacks the conserved aromatic residue in the F strand between Cys3 and Cys4, and DMBT1 lacks the second conserved aromatic residue after Cys4. These two residues interact in the ZP3-N crystal structure (Monne et al., 2008) and are thought to be critical for ZPn polymerization. Genbank accession numbers: Ce_LET-653b: NP_001021337; Ce_DYF-7: NP_509630; Dm_Dpy: NP_001245875; Hs_UMOD: NP_003352; Hs_TectA: NP_005413; Hs_ZP3: NP_001103824; Hs_DMBT1: NP_004397; Hs_Oit3: NP_689848; Hs_Eng: NP_001108225.

**Table S1.** Strains generated in this study.

| Strain | Genotype | Citations |
| --- | --- | --- |
| UP3121 | *csEx624 [let-653pro::SfGFP::LET-653(ZP); lin48pro::mRFP]* | (Gill et al., 2016) |
| UP3244 | *let-653(cs178) IV; csEx358 [lpr-1pro::LET-653b; unc-119pro::GFP]* | (Gill et al., 2016) |
| UP3342 | *let-653(cs178) IV; csEx766 [lin48pro::LET-653b::SfGFP; myo-2pro::GFP]* | (Forman-Rubinsky et al., 2017) |
| UP3422 | *csIs66 [let-653pro::LET-653(ZP)::SfGFP; let-653pro::PH::mCherry] X* | (Forman-Rubinsky et al., 2017) |
| UP3432 | *let-653(cs178) IV; csEx821 [let-653pro::SfGFP::LET-653(ZPc); lin-48pro::mRFP]* |  |
| UP3448 | *csEx822 [let-653pro::SfGFP::LET-653(ZPc); lin-48pro::mRFP]* |  |
| UP3449 | *csEx821 [let-653pro::SfGFP::LET-653(ZPc); lin-48pro::mRFP]* |  |
| UP3462 | *let-653(cs178) IV; csIs66 [let-653pro::LET-653(ZP)::SfGFP; let-653pro::PH::mCherry] X* |  |
| UP3465 | *let-653(cs178) IV; csEx828 [let-653pro::LET-653(ZPc); lin-48pro::mRFP]* |  |
| UP3466 | *let-653(cs178) IV; csEx829 [let-653pro::LET-653(ZPc); lin-48pro::mRFP]* |  |
| UP3514 | *csEx841 [let-653pro::LET-653(ZPc)::SfGFP; lin-48pro::mRFP]* |  |
| UP3515 | *csEx842 [let-653pro::LET-653(ZPc)::SfGFP; lin-48pro::mRFP]* |  |
| UP3516 | *csEx843 [let-653pro::SfGFP::LET-653(ZPn); lin-48pro::mRFP]* |  |
| UP3517 | *csEx844 [let-653pro::SfGFP::LET-653(ZPn); lin-48pro::mRFP]* |  |
| UP3594 | *let-653(cs178) IV; csEx841 [let-653pro::LET-653(ZPc)::SfGFP; lin-48pro::mRFP]* |  |
| UP3630 | *csEx882 [let-653pro::LET-653(ZP,AYAA)::SfGFP; lin-48pro::mRFP]* |  |
| UP3630 | *csEx882 [let-653pro::LET-653(ZP,AYAA)::SfGFP; lin-48pro::mRFP]* |  |
| UP3631 | *csEx883 [let-653pro::LET-653(ZP,AYAA)::SfGFP; lin-48pro::mRFP]* |  |
| UP3746 | *let-653(cs262 [LET-653::SfGFP]) IV* | (Cohen et al., 2020) |
| UP3777 | *let-653(cs178) IV; csEx889 [let-653pro::SfGFP::LET-653(ZPn); lin-48pro::mRFP]; csEx358* |  |
| UP3806 | *let-653(cs178) IV; csEx885 [let-653pro::LET-653(ZP,AYAA); lin-48pro::mRFP]* |  |
| UP3807 | *let-653(cs178) IV; csEx893 [let-653pro::SfGFP::LET-653(ZPn); lin-48pro::mRFP]; csEx358* |  |
| UP3843 | *csIs91 [let-653pro::SfGFP::LET-653(ZPc); lin-48pro::mRFP]* |  |
| UP3844 | *csIs92 [let-653pro::SfGFP::LET-653(ZPc);* |  |
|  | *lin-48pro::mRFP* |  |
| UP3845 | *csIs93 [let-653pro::SfGFP::LET-653(ZPc);*<br>*lin-48pro::mRFP]* |  |
| UP3846 | *csIs94 [let-653pro::LET-653(ZP, AYAA)::SfGFP;*<br>*lin-48pro::mRFP]* |  |
| UP3847 | *csIs95 [let-653pro::LET-653(ZP, AYAA)::SfGFP;*<br>*lin-48pro::mRFP]* |  |
| UP3848 | *csEx905 [let-653pro::LET-653(ZP)::SfGFP;*<br>*lin-48pro::mRFP]* |  |
| UP3849 | *csIs96 [let-653pro::LET-653(ZP)::SfGFP;*<br>*lin-48pro::mRFP]* |  |
| UP3865 | *csEx913 [let-653pro::SfGFP::LET-653(ZPc-½Cterm);*<br>*lin-48pro::mRFP]* |  |
| UP3866 | *csEx916 [let-653pro::LET-653(ZPn+Cterm)::SfGFP;*<br>*lin-48pro::mRFP]* |  |
| UP3867 | *csEx912 [let-653pro::SfGFP::LET-653(ZPc-½Cterm);*<br>*lin-48pro::mRFP]* |  |
| UP3868 | *csEx911 [let-653pro::SfGFP::LET-653(ZPc-½Cterm);*<br>*lin-48pro::mRFP]* |  |
| UP3869 | *csEx910 [let-653pro::SfGFP::LET-653(ZPc-½Cterm);*<br>*lin-48pro::mRFP]* |  |
| UP3870 | *csEx917 [let-653pro::LET-653(ZPn+Cterm)::SfGFP;*<br>*lin-48pro::mRFP]* |  |
| UP3871 | *csEx915 [let-653pro::LET-653(ZPn+Cterm)::SfGFP;*<br>*lin-48pro::mRFP]* |  |
| UP3872 | *csEx914 [let-653pro::LET-653(ZPn+Cterm)::SfGFP;*<br>*lin-48pro::mRFP]* |  |
| UP3880 | *let-653(cs178) IV; csEx696 [let-653pro::*<br>*LET-653(ZP)::SfGFP; lin-48pro::mRFP]* |  |
| UP3919 | *let-653(cs178) IV; csEx913 [let-653pro::SfGFP::*<br>*LET-653(ZPc-½Cterm); lin-48pro::mRFP]; csEx766* |  |
| UP3929 | *let-653(cs178) IV; csEx915 [let-653pro::*<br>*LET-653(ZPn+Cterm)::SfGFP; lin-48pro::mRFP]; csEx358* |  |
| UP3930 | *let-653(cs178) IV; csEx917 [let-653pro::*<br>*LET-653(ZPn+Cterm)::SfGFP; lin-48pro::mRFP]; csEx358* |  |
| UP3931 | *let-653(cs178) IV; csEx915 [let-653pro::*<br>*LET-653(ZPn+Cterm)::SfGFP; lin-48pro::mRFP]; csEx358* |  |
| UP3951 | *let-653(cs178) IV; csEx911 [let-653pro::SfGFP::*<br>*LET-653(ZPc-½Cterm); lin-48pro::mRFP]; csEx766* |  |
| UP3952 | *let-653(cs178) IV; csEx886 [let-653pro::*<br>*LET-653(ZP,AYAA)::SfGFP; lin-48pro::mRFP]; csEx358* |  |
| UP3953 | *let-653(cs178) IV; csIs96 [let-653pro::*<br>*LET-653(ZP)::SfGFP; lin-48pro::mRFP]* |  |
| UP3988 | *csEx934 [let-653pro::LET-653(ZPc,AYAA)::SfGFP;*<br>*lin-48pro::mRFP]* |  |
| UP3997 | *csEx935 [let-653pro::LET-653(ZPc,AYAA)::SfGFP;*<br>*lin-48pro::mRFP]* |  |
| UP3998 | *csEx936 [let-653pro::LET-653(ZPc,AYAA)::SfGFP;*<br>*lin-48pro::mRFP]* |  |
| UP3999 | *let-653(cs178) IV; csEx938 [let-653pro::*<br>*LET-653(ZPc,AYAA)::SfGFP; lin-48pro::mRFP]; csEx358* |  |
| UP4000 | *let-653(cs178) IV; csEx937 [let-653pro::*<br>*LET-653(ZPc,AYAA)::SfGFP; lin-48pro::mRFP]; csEx358* |  |

**Table S2.**
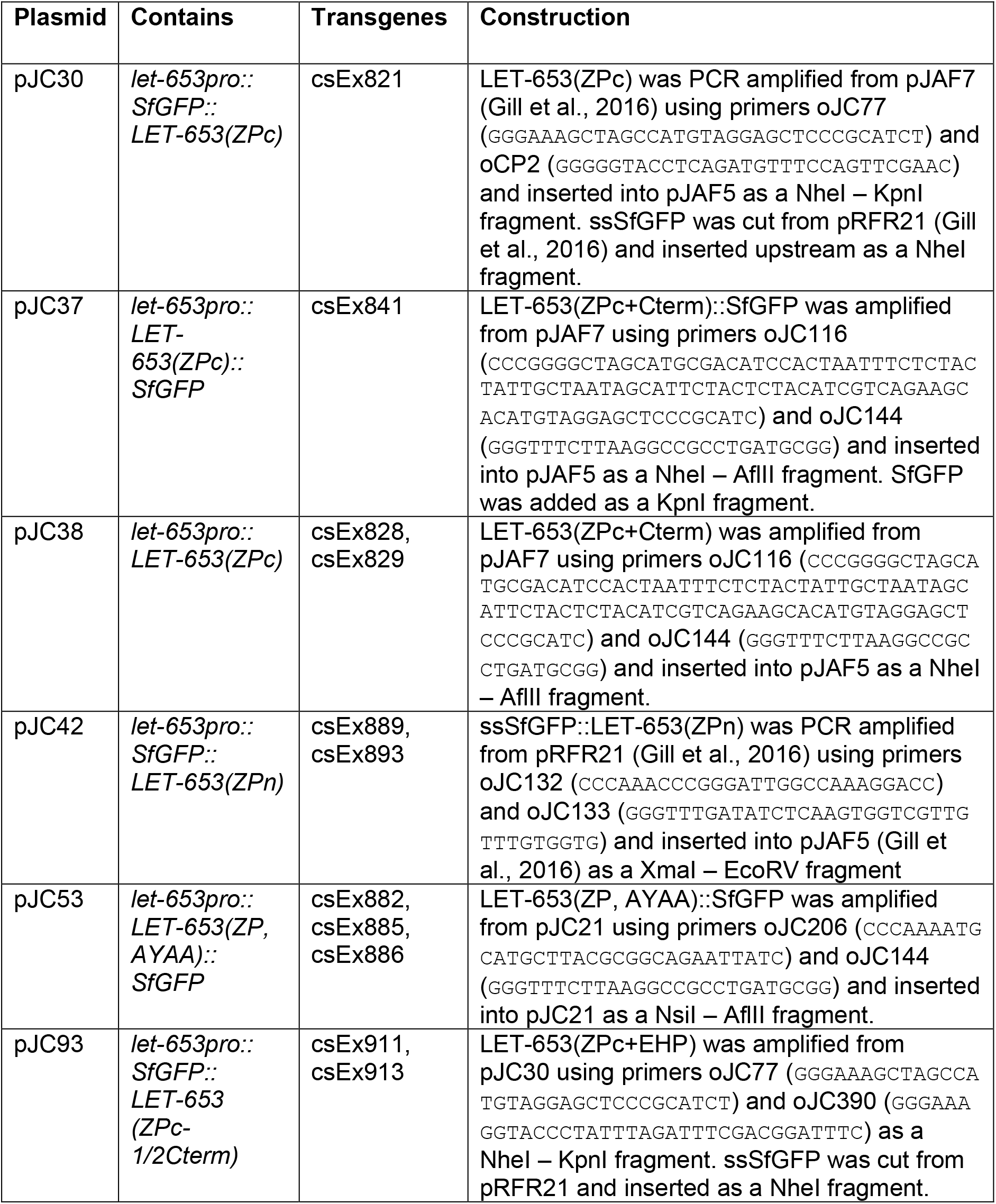
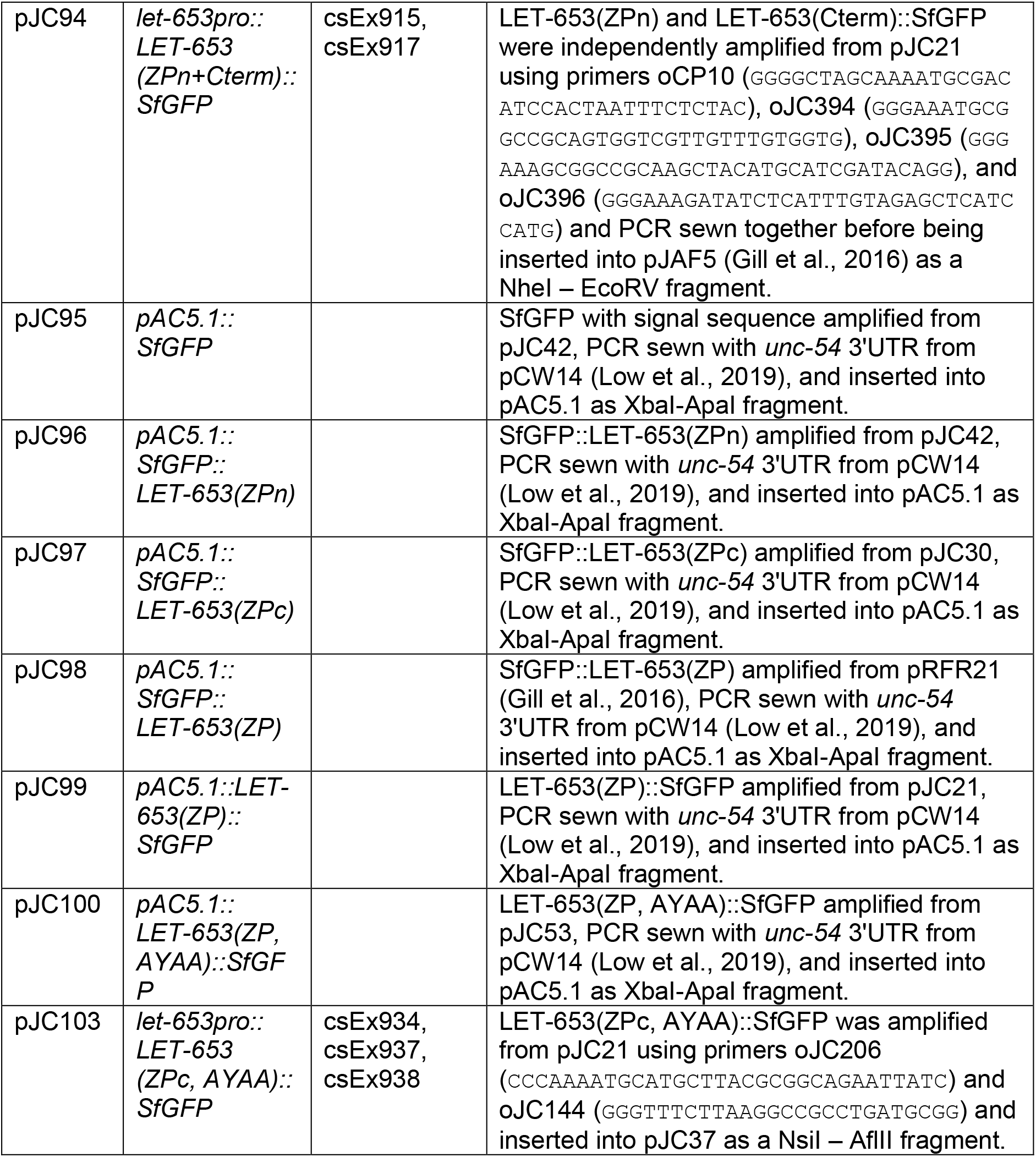
Plasmids generated in this study

